# Cardiac Timing Biases Creative Exploration and Exploitation

**DOI:** 10.64898/2026.06.03.729883

**Authors:** Francesca Torno Jimenez, James Lloyd-Cox, Caroline Di Bernardi Luft, Maria Herrojo Ruiz, Joydeep Bhattacharya

## Abstract

Creative ideation involves a dynamic exchange between exploring new idea categories and exploiting familiar ones, reflecting optimal foraging principles. Although interoceptive signals, particularly cardiac activity, are associated with differences in attention and cognitive control, their role in explore-exploit dynamics during creative idea generation remains unknown. Recording both electroencephalography (EEG) and electrocardiography (ECG) data, we used convolution general linear models to examine cardiac-brain interactions underlying semantic exploration and exploitation during creative idea generation, in both spontaneous (self-selected) and directed (externally cued) conditions. Cardiac deceleration predicted ideation time: this relationship scaled nonlinearly during exploration (category switching), but linearly during exploitation (category persistence), in both conditions. Cardiac deceleration did not predict semantic distance (between response and cue word) or response accuracy during the directed condition, suggesting that heart rate slowing reflects cognitive effort allocation rather than ideational content. Cardiac phase systematically biased explore-exploit dynamics: spontaneous switching and accurate directed-switching were preferentially associated with diastolic phases and parieto-occipital alpha (8-12 Hz) desynchronization, whereas category persistence was associated with systolic phase timing and frontal theta (4-6 Hz) synchronization. The former activity was coupled to higher CD, as expected. However, the latter time-frequency activity was linked to responses timed to systole. These findings suggest that creative ideation unfolds through embodied cardiac-cortical coordination, whereby diastolic states are associated with flexible semantic exploration and alpha-related attentional dynamics, whereas systolic states are associated with exploitative persistence and theta-related control processes. This suggests interoceptive rhythms as temporal scaffolding that structures when and how ideas emerge, fundamentally expanding creativity neuroscience beyond purely cortical models.

## Introduction

Human creativity depends on the capacity to dynamically navigate semantic memory, alternating between exploitation of familiar conceptual terrain and exploration of distant, novel associations (Kenett & Faust, 2019). When humans generate creative ideas, they alternate between persistent search within semantic categories (e.g., listing foods) and flexible shifts to new conceptual regions (e.g., switching from “food” to “occupations”). This dynamic balance between local exploitation and global exploration has long been recognized as central to optimal foraging in animals navigating physical environments (Charnov, 1976) yet appears equally fundamental to internal search through memory (Hills et al., 2015; Malaie et al., 2024; Todd & Hills, 2020). Despite substantial progress in characterizing the structure of semantic search in creative cognition (Kenett & Beaty, 2023; Hart et al., 2017), the neurocognitive mechanisms that govern when and how individuals transition between semantic exploration and exploitation remain incompletely understood. While extensive research has characterized the cortical networks supporting creative ideation (Beaty et al., 2016; Benedek & Jauk, 2019), it remains unknown whether interoceptive signals, particularly those linked to cardiac activity, contribute to shifts between exploration and exploitation. This possibility is consistent with the anatomical and functional overlap between creativity-related control networks and systems involved in monitoring bodily states (Craig, 2009; Critchley et al., 2004). This study addresses this gap by examining the neurophysiology of cardiac influences on exploration and exploitation strategies during creative ideation, both when these strategies emerge spontaneously and when they are externally imposed.

Creativity is traditionally measured through divergent thinking tasks (Silvia et al., 2008; Acar & Runco, 2019) such as the Alternate Uses Task (AUT; Guilford, 1967), which requires participants to generate as many creative uses for common objects as possible (e.g. using a brick to draw on a chalkboard). Responses are typically assessed for fluency (quantity of responses), flexibility (number of semantic categories covered), originality (statistical rareness of the responses), and elaboration (extent to which responses are described and embellished; Reiter-Palmon et al., 2019). Performing divergent thinking tasks like the AUT requires the ability to search within semantic memory and generate ideas outside common associations (Kenett & Faust, 2019; Hass, 2017; Ovando-Tellez et al., 2022; Beaty et al., 2021). After generating a creative use, participants can follow one of two strategies: either exploiting the current semantic category in subsequent responses, finding similar creative uses (e.g. brick → to make a mark → to make a stamp), or exploring new semantic categories (e.g. brick → to make a mark → to grind up food). These strategies are selected locally based on the previous idea and change depending on the individual’s estimate of when a semantic space is depleted (Hart et al., 2017; Hills et al., 2015). As ideation proceeds, responses tend to become more creative and more remote from initial ideas (Beaty & Silvia, 2012; Wang et al., 2017). These patterns mirror optimal foraging behavior in animals: periods of local exploitation (area-restricted search) alternating with exploratory jumps to new resource areas, with transitions occurring when local retrieval rates decline below the expected global average (Charnov, 1976; Todd & Hills, 2020). Computational models also support the notion that humans navigate semantic space through a combination of similarity-guided local exploitation and frequency-guided global exploration, alternating between high-clustering “persist” periods and long-distance “switch” events (Hills et al., 2012; Kumar, 2020; Lundin et al., 2023).

Individual differences in the ability to shift between exploiting familiar categories and exploring new semantic spaces can greatly affect divergent thinking performance (Baror & Bar, 2016; Hart et al., 2017; Vartanian et al., 2007; Koutstaal, 2025). Less creative individuals often exhibit stronger connections between common associates in semantic memory (e.g., brick → home) and weaker connections between uncommon associates (e.g., brick → stamp), whereas highly creative individuals exhibit more balanced strength between both common and uncommon associations (Benedek et al., 2017; Kenett et al., 2014). Such differences in semantic memory structure may underlie individual differences in the ability to shift between exploratory and exploitative strategies. Indeed, this ability to shift between strategies has also been linked to associative and controlled memory processes (Beaty et al., 2021; Benedek & Jauk, 2019; Benedek et al., 2023), where associative processes promote exploration and controlled processes promote exploitation (Nijstad et al., 2010). Understanding the mechanisms that drive these transitions between exploratory (semantic switching) and exploitative (semantic persistence) remains a central question in the study of human cognition.

### Neural systems supporting creativity

Exploration and exploitation strategies are driven by two large-scale brain networks: the executive control network (ECN), supporting goal-directed control and mental set shifting, and the default mode network (DMN), supporting spontaneous associative ideation (Bartoli et al., 2024; Bashwiner et al., 2016; Beaty et al., 2014, 2015, 2016; Dietrich, 2004; Gabora, 2018; Lloyd-Cox et al., 2025; Rosen et al., 2020; Shi et al., 2017; Wise & Braga, 2014; but also see Abraham, 2025). These networks are typically anti-correlated – when one activates, the other deactivates (Goulden et al., 2014) – yet increased connectivity between DMN and ECN regions predicts enhanced performance on verbal divergent thinking (Beaty et al., 2015; Green et al., 2015). Moreover, the frequency of switching between DMN and ECN at rest has been linked to divergent thinking ability (Chen et al., 2025). Critically, switching between these networks is orchestrated by a third network, the salience network, SN (Goulden et al., 2014; Menon & Uddin, 2010). Regions of the SN, including the anterior cingulate cortex (ACC) and the insular cortices (IC), have been linked to creative cognition and are proposed to bring different types of thinking to awareness depending on task demands (Beaty et al., 2015; Chrysikou et al., 2014; Heinonen et al., 2016; Huo et al., 2025). These same regions are also the primary cortical hubs for processing interoceptive stimuli, dynamically influencing the allocation of cognitive resources based on bodily signals (Craig, 2009; Ceunen et al., 2016; Chong et al., 2017; Critchley et al., 2004; García-Cordero et al., 2017; Haruki & Ogawa, 2021; Pollatos et al., 2005; Zaki et al., 2012). This anatomical overlap raises a fundamental question: if the same neural regions underlie both creative ideation strategies and interoceptive awareness, do bodily signals, particularly cardiac rhythms, contribute to the transition between exploratory and exploitative semantic search? While much is known about the neural systems supporting divergent thinking, the influence of physiological signals on creative ideation remains largely unknown.

Electroencephalographic (EEG) studies have identified frequency-specific oscillatory signatures of creative cognition. Frontal alpha (8-12 Hz) synchronization, indicative of cortical inhibition (Klimesch et al., 2007), is consistently associated with greater creative performance (Benedek et al., 2011; Camarda et al., 2018; Fink et al., 2018; Fink & Benedek, 2014; Luft et al., 2018; Stevens & Zabelina, 2020). Heightened alpha activity, especially in temporo-parietal regions, may reflect reduced external sensory input, allowing greater attention to internal cognitive processes during divergent thinking (Benedek et al., 2016; Luft et al., 2018; Zioga et al., 2024). Beyond alpha, mid-frontal theta (4-7 Hz) oscillations, often associated with cognitive control (Cavanagh & Frank, 2014), are also implicated in creative ideation (Bartoli et al., 2024). These cortical oscillations, however, have been studied almost exclusively in isolation from interoceptive signals that might rhythmically structure when and how ideas are generated.

### Cardiac signals that modulate cognition

Emerging evidence suggests that cardiac signals modulate both attention and the cortical oscillations implicated in creative cognition. Heightened cardiac signaling to the brain (e.g. during baroreceptor activation and other states of cardiovascular arousal) selectively attenuates attention to external stimuli in favor of internally directed attention (Al et al., 2020; Garfinkel et al., 2014; Ren et al., 2022, 2024), raising the possibility that cardiac signals may influence the gating of sensory input during creative ideation. This is consistent with evidence that heightened alpha activity, especially in temporo-parietal regions, reflects reduced external sensory input, allowing greater attention to internal cognitive processes during divergent thinking (Benedek et al., 2016; Zioga et al., 2024; Luft et al., 2018). Cardiac activity has also been shown to modulate alpha power (Kawashima et al., 2024; Lai et al., 2024; Luft & Bhattacharya, 2015; Magosso et al., 2019), suggesting that heartbeat signals may shape the oscillatory states that support creative ideation (Baas et al., 2008). Despite these suggestive findings, past research linking cardiac signals to creative performance remains mixed, and the underlying mechanisms are poorly understood (Loudon & Deininger, 2016; Rominger et al., 2022; Silvia et al., 2014).

For instance, Silvia et al. (2014) found that individuals reporting greater creative achievement exhibit increases in AUT performance associated with sympathetic activity, but no relationship with parasympathetic activity as measured through heart rate variability (HRV). Rominger and colleagues (2022) found that HRV is significantly tied to creative thinking, with this relationship moderated by physical activity levels: individuals who exercised regularly experienced HRV increases during divergent thinking, while those who did not exercise showed HRV decreases. Earlier research by Loudon and Deininger (2016) found that decreases in low-frequency HRV (i.e. lower sympathetic-related activity) were associated with greater fluency on divergent thinking tasks. Collectively, while these findings focus on changes in HRV, the specific mechanisms that link cardiac dynamics to ideational strategies – particularly the balance between exploration and exploitation – remain unexplored. No prior study has examined whether cardiac timing systematically biases when people persist within versus switch between semantic categories during creative ideation, or how such timing interacts with cortical activity. Understanding these processes, including changes in heart rate and the timing of the cardiac cycle, is critical to understanding how humans balance explore-exploit dynamics during creative ideation.

The baroreceptor hypothesis provides a mechanistic framework for understanding cardiac influences on cognition. The hypothesis posits that the cardiovascular system affects central cortical excitability via changes in heart rate and blood pressure (Lacey & Lacey, 1978; Elbert & Rau, 1995), responding to environmental demands and bodily energy requirements (Duschek et al., 2013). Cardiac cycles consist of systole (ventricular contraction ejecting blood into the aorta) and diastole (ventricular relaxation, during which the atria refill with oxygenated blood). These cycles are influenced by parasympathetic and sympathetic activity to accelerate or decelerate heart rate depending on environmental demands (Graham & Clifton, 1966). During systole, baroreceptors (i.e., nerve cells located in the aortic arch and carotid sinus that detect blood pressure changes by sensing the stretching of vessel walls) signal elevated arterial pressure to brainstem nuclei, which relay this information to the insula and ACC, potentially modulating cortical activity by inhibiting sensory processes (Al et al., 2020; Elbert & Rau, 1995). This may explain why people exhibit reduced sensitivity to sensory stimuli when it is delivered during systole (Elbert & Rau, 1995; Park & Blanke, 2019; Pramme et al., 2014, 2016). For instance, electrocutaneous pain administered during systole is perceived with less intensity than if it is administered during diastole (Droste et al., 1994; Wilkinson et al., 2013).

However, systole-related effects on cognition are mixed and appear to depend on task demands. While systolic presentation reduces pain sensitivity (Droste et al., 1994; Wilkinson et al., 2013), it can also enhance responsiveness to emotionally salient stimuli (Adelhöfer et al., 2020; Garfinkel et al., 2014), indicating that its impact varies depending on the context. Additionally, research indicates that systole increases the likelihood of spontaneous (self-initiated) exploratory behaviors (Kunzendorf et al., 2019; Palser et al., 2021), which may reflect an advantageous disposition for spontaneous exploration in creative contexts. The heartbeat-evoked potential (HEP), a cortical electrophysiological response time-locked to each heartbeat, provides a neural marker of cardiac-cortical coupling (Coll et al., 2021): higher pre-stimulus HEP amplitude predicts reduced perceptual detection of near-threshold stimuli (Park et al., 2014; Skora et al., 2022), suggesting that ongoing cardiac signaling competes with exteroceptive processing. Cardiac deceleration, a parasympathetic marker reflecting adaptive allocation of cognitive resources, accompanies increased cognitive workload and sustained attention (Hashemi et al., 2019; Roelofs et al., 2010; Van Roon et al., 2004), suggesting that heart rate slowing may index the computational demands of semantic exploration. Yet whether and how these cardiac dynamics influence transitions between semantic switching and semantic persistence during creative ideation remains unknown.

### The current study

Three critical gaps emerge from this literature. First, although cardiac phase effects have been documented across multiple cognitive domains, their role in regulating exploration and exploitation within semantic memory remains unknown. This omission is consequential, given that theories of brain-body interaction predict that systolic and diastolic timing may differentially bias categorical switching versus category persistence. Second, although cardiac signals modulate the cortical rhythms central to creativity, particularly alpha and theta, no study has investigated whether cardiac-oscillatory coupling is reflected in the neural signatures of switching and persistence during idea generation. Third, existing creativity research has focused almost exclusively on cortical mechanisms, ignoring the continuous interoceptive inputs to the brain that might rhythmically structure when ideas emerge. This represents a fundamental conceptual gap: if creative cognition involves dynamic coordination by the salience network, a system fundamentally organized around interoceptive signals, then creativity may be inherently embodied, with cardiac rhythms serving as temporal scaffolding for ideational transitions.

This study provides the first systematic examination of how cardiac dynamics relate to creative behaviors and cortical oscillatory activity, linking physiological signals with the regulation of exploration and exploitation in human cognition. We aimed to clarify the role of cardiac activity in facilitating shifts between exploratory and exploitative creative search strategies, providing the first embodied account of creativity at the level of cardiac-neural dynamics. By demonstrating that cardiac timing systematically biases creative behaviors, this work fundamentally expands neuroscientific understanding of creative cognition beyond purely cortical models to encompass the body’s continuous rhythmic influence on the temporal dynamics of creativity.

We tested three central hypotheses grounded in the integration of interoception theory, baroreceptor physiology, and creativity neuroscience. First, drawing on models where cardiac deceleration reflects cognitive effort allocation (Hashemi et al., 2019; Roelofs et al., 2010; Van Roon et al., 2004), and given that semantic exploration is computationally demanding and associated with longer response times (Kenett & Faust, 2019; Mastria et al., 2021; Mazza et al., 2023), we predicted that greater cardiac deceleration would precede longer response times and an increased likelihood of spontaneous semantic switching. Second, based on baroreceptor models showing systolic inhibition of cortical excitability and diastolic facilitation of information processing (Edwards et al., 2007; Motyka et al., 2019; Fiacconi et al., 2016), we predicted that spontaneous semantic switching would preferentially occur during systole – when reduced external sensory interference may facilitate internally directed cognition – and that semantic persistence would preferentially occur during diastole, particularly under high control demands. Third, we predicted that these cardiac–behavioral interactions would be mirrored in oscillatory brain activity patterns previously implicated in creative cognition and executive control, particularly in the alpha band (8–12 Hz) for semantic search and internal focus (Fink & Benedek, 2014; Jauk et al., 2012), and theta band (4–6 Hz) for mental set maintenance and top-down control (Cavanagh & Frank, 2014; Luft, 2014). This integrative approach allows us to establish whether ideational shifts during creative cognition are governed not solely by cortical dynamics, but also temporally aligned with cardiac signaling, positioning creative cognition as an inherently embodied process.

## Results

Participants (*N* = 59) completed two versions of the Alternate Uses Task (AUT) while we recorded concurrent EEG and ECG (see Fig. 1 for the experimental design). In the spontaneous (AUT; classical version) condition, participants generated creative uses for common objects (e.g., brick, drinking straw) while alternating freely between semantic exploration (switching to new idea categories) and semantic exploitation (persisting within a single category). In the directed condition (D-AUT), participants received explicit instructions (randomly selected, while maintaining an equal number of switches and persists) after each response: an ‘S’ indicated that the next idea should be similar to their previous idea (i.e., they should persist within the same semantic category; directed-persist condition); a ‘D’ indicated that the next idea should be different to their previous idea (i.e., they should switch to a new semantic category; directed-switch condition). This within-subject manipulation allowed assessment of cardiac-brain dynamics under spontaneous and externally instructed ideation behaviors.

**Figure 1.**
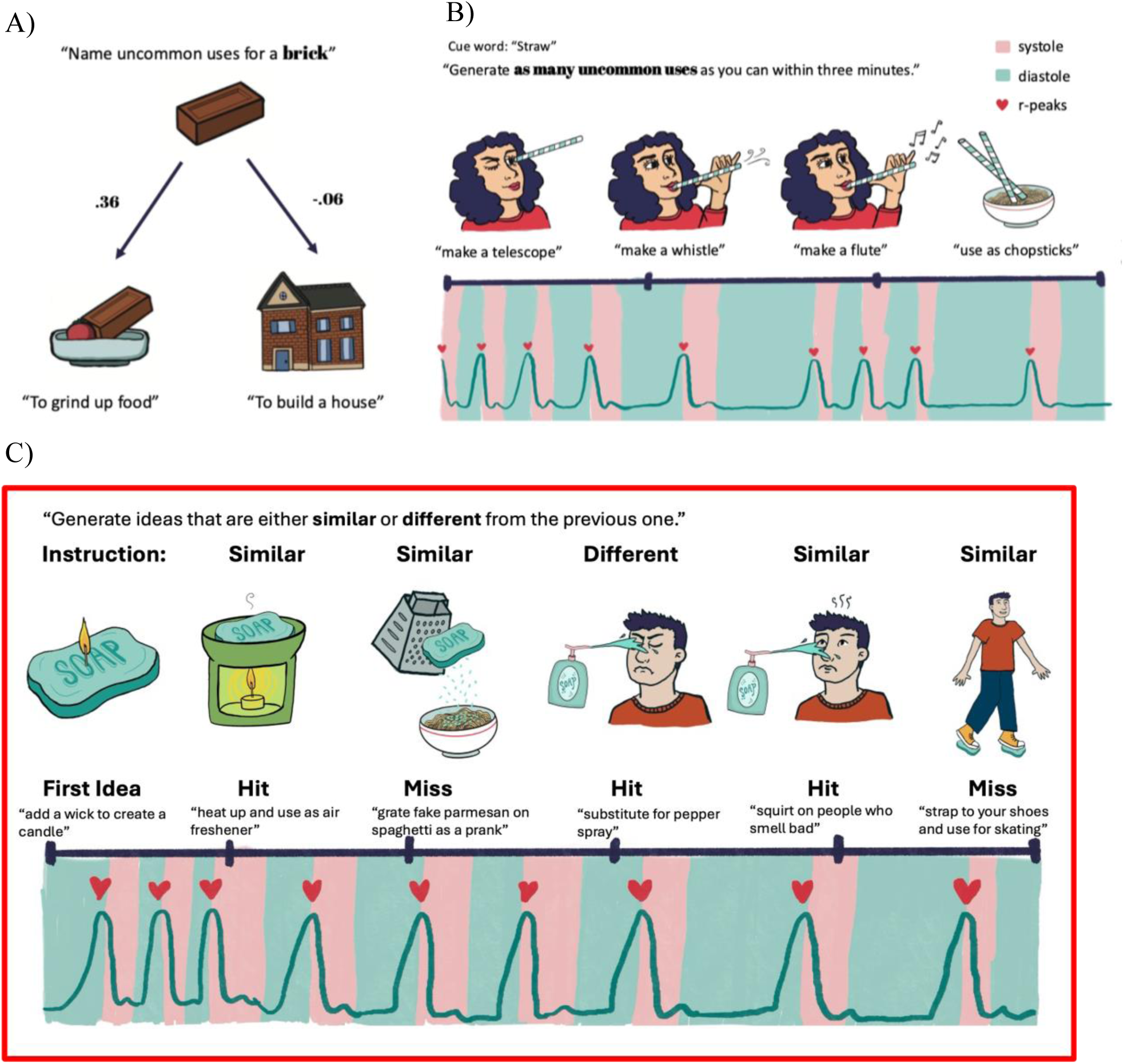
Experimental design and measurement approach. A) Semantic distance between a cue word (“brick”) and example participant responses. **B)** Schematic of the spontaneous Alternate Uses Task (AUT) with concurrent EEG and ECG recording. Responses shown are real participant examples. **C)** Schematic of the directed AUT (D-AUT), illustrating examples of accurate semantic switches and accurate semantic persistence. Responses shown are real participant examples.

For each trial (instance of idea generation), we extracted four key variables: (i) inter-response semantic distance; the semantic distance between consecutive responses estimated using the SemDis model (Beaty & Johnson, 2021), (ii) response time, indexed by the inter-response interval, (iii) cardiac deceleration (CD), quantified as the change in heart rate from pre-response baseline (including three inter-beat intervals before, one during/after, and two after button-press), and (iv) cardiac phase at the moment of button-press (systole vs. diastole). To address our questions regarding the relationship between brain rhythms and creative behaviors, EEG data were decomposed into time-frequency representations spanning the 4-30 Hz range, thus including three frequency bands: theta (4-6 Hz), alpha (8-12 Hz), and beta (13-30 Hz). We then modelled changes in EEG amplitudes in this time-frequency representation using a convolution GLM (Litvak et al., 2013). Convolution GLMs allow modelling the effect of one regressor on the neural signals, while controlling for the effect of other regressors included in the model (like standard GLMs). The advantage of convolution GLM is that it can robustly estimate neural changes to regressors when events have overlapping and varying latencies trial to trial (Litvak et al., 2014). This was relevant in our case, where we modelled pseudocontinuous time series by concatenating long epochs from the moments leading up to the ideation-related button-press (-3 to + 2 s around event onset).

We used two separate convolution GLMs for the AUT and the D-AUT. The AUT convolution model included two regressors: one parametric regressor (inter-response semantic distance, or SemDis) and one discrete regressor (button_press). For the D-AUT, five discrete regressors were used (switch_hit, switch_miss, persist_hit, persist_miss, and button_press). Continuous regressor SemDis was chosen for the GLM of the AUT given that spontaneous ideation cannot be univocally split into binary categories as for the D-AUT. For the AUT, this approach allowed us to model linear relationship in alpha and theta rhythms to the breadth of semantic distances covered by participants’ responses. For the D-AUT, this approach allowed us to model within-person differences in alpha and theta rhythms to accuracy on both switching and persisting, rather than the semantic breadth of responses. For both conditions, using button-press as a regressor allowed us to control for motor activity which might have otherwise been conflated with the behavior of interest. Moreover, we also averaged GLM outputs for analyzing the relationship between brain-behavior activity and cardiac signals.

To answer our questions, we employed four complementary levels of analysis: (i) Individual differences to identify relationships between stable cardiac signals (i.e.: heart rate, cardiac phase ratio) on ideation tendencies (i.e.: semantic distance, response times), (ii) Trial-level mixed-effects models to capture moment-to-moment cardiac-behavior coupling, (iii) Simple linear regressions linking cardiac measures with averaged GLM outputs of time-frequency responses, and (iv) Circular statistics to model the cyclical nature of cardiac phases. Finally, we conducted control analyses related to possible motor confounds in the beta frequency band as well as exploratory analyses related to the originality of responses (semantic distance from the cue word to the response, in contrast to the inter-response semantic distance used to assess switching).

### Behavioral performance across conditions

Participants demonstrated normal engagement across all task conditions (see Table 1 and Supplementary Table 1 for descriptive statistics and ANOVAs of condition-related differences). For the AUT, participants generated an average of 29.6 ideas (SD = 12.1, range = 8–68) across 2,947 valid trials. The directed-switch and directed-persist (D-AUT) conditions yielded comparable outputs, with means of 31.1 (SD = 14.4, range = 7–75; 1,759 trials) and 32.5 (SD = 13.5, range = 9–82; 1,812 trials), respectively.

**Table 1.** Descriptive statistics for main variables by condition. This table presents descriptive statistics and results from repeated-measures ANOVAs comparing behavioral and physiological measures across three conditions: spontaneous ideation, directed-persist (exploitative), and directed-switch (exploratory). Semantic Distance differed significantly across conditions, *F*(2, 174) = 20.64, *p* < .001, *η²* = .192. The directed-persist condition exhibited lower semantic distance values relative to the spontaneous and directed-switch conditions, consistent with reduced exploratory (i.e., semantically distant) responses during persistence. Response Time did not significantly vary across conditions, *F*(2, 174) = 2.04, *p* = .133, *η²* = .023, suggesting that differences in ideation strategy were not driven by changes in response speed. For physiological measures, Cardiac Phase Ratio showed a marginal effect of condition, *F*(2, 174) = 2.70, *p* = .07, *η²* = .030, indicating a potential trend toward differences in the timing of responses within the cardiac cycle. However, Inter-beat Interval (IBI) did not differ across conditions, *F*(2, 174) = 0.46, *p* = .633, *η²* = .005, suggesting overall stable heart rate across task demands.

| Variable | Spontaneous<br>M (SD) | Directed-Persist<br>M (SD) | Directed-Switch<br>M (SD) | F(df1, df2) | *p* | $\eta^2$ |
| --- | --- | --- | --- | --- | --- | --- |
| Ideation Strategy<br>Ratio | 0.79 (0.08) | 0.42 (0.17) | 0.85 (0.10) | 217.13 (2, 174) | < .001 | 0.714 |
| Semantic Distance | 0.01 (0.02) | -0.02 (0.04) | 0.01 (0.02) | 20.64 (2, 174) | < .001 | 0.192 |
| Response Time | 14.98 (4.91) | 13.20 (4.65) | 14.21 (4.87) | 2.04 (2, 174) | 0.133 | 0.023 |
| Cardiac Phase<br>Ratio | 0.40 (0.10) | 0.37 (0.15) | 0.41 (0.10) | 2.70 (2, 174) | 0.07 | 0.030 |
| Inter-beat interval<br>(IBI) | 0.80 (0.09) | 0.81 (0.09) | 0.81 (0.09) | 0.46 (2, 174) | 0.633 | 0.005 |

### Cardiac deceleration (CD) predicts response time

We next investigated our first hypothesis, whether greater CD predicted two key behavioral metrics: inter-response semantic distance and response time, using mixed-effects models. Full model specifications and estimates are reported in Table 2. Contrary to our expectations, CD showed no significant relationship with inter-response semantic distance (*ps* > .100) in any of the conditions.

**Table 2.** Table of mixed effects models examining the relationship between SemDis and Cardiac Phase. Model numbers reflect model structure (Model 1: RespPhaseNumeric; Model 2: RespPhaseNumeric + beforeRespIBI; Model 3: RespPhaseNumeric × beforeRespIBI), whereas letters denote experimental condition (A = spontaneous, B = directed persist, C = directed switch). All models include random intercepts for participants (Pnum) and trials (IDinTask). These linear mixed-effects models were run using two fixed factors, cardiac phase (RespPhaseNumeric – where 1 = systole, 0 = diastole) and average inter-beat interval lengths leading up to response (beforeRespIBI), semantic distance of the response to the response prior (SemDis) as an outcome.

| Predictor | Estimate<br>($\beta$) | SE | df | t | p | Model |
| --- | --- | --- | --- | --- | --- | --- |
| **Model 1A:** SemDis ~ RespPhaseNumeric + (1 \| Pnum) + (1 \| IDinTask) |  |  |  |  |  |  |
| Intercept | .01 | .00 | 97.20 | 3.04 | .00 | 1A |
| RespPhaseNumeric | .00 | .00 | 2407.00 | .47 | .64 | 1A |
| **Model 2A:** SemDis ~ RespPhaseNumeric + beforeRespIBI + (1 \| Pnum) + (1 \| IDinTask) |  |  |  |  |  |  |
| Intercept | .00 | .02 | 290.30 | .26 | .80 | 2A |
| RespPhaseNumeric | .00 | .00 | 2405.00 | .48 | .63 | 2A |
| beforeRespIBI (s) | .01 | .02 | 297.00 | .27 | .79 | 2A |
| **Model 3A:** SemDis ~ RespPhaseNumeric × beforeRespIBI + (1 \| Pnum) + (1 \| IDinTask) | | | | | | |
| Intercept | .01 | .02 | 516.00 | .56 | .58 | 3A |
| RespPhaseNumeric | -.015 | .03 | 2396.00 | -.57 | .57 | 3A |
| beforeRespIBI (s) | -.003 | .03 | 532.00 | -.11 | .91 | 3A |
| RespPhaseNumeric × beforeRespIBI | .02 | .03 | 2396.00 | .63 | .53 | 3A |
| **Model 1B:** SemDis ~ RespPhaseNumeric + (1 \| Pnum) + (1 \| IDinTask) |  |  |  |  |  |  |
| Intercept | -.013 | .01 | 72.70 | -2.09 | .04 | 1B |
| RespPhaseNumeric | -.017 | .01 | 1240.00 | -2.23 | .03 | 1B* |
| **Model 2B:** SemDis ~ RespPhaseNumeric + beforeRespIBI + (1 \| Pnum) + (1 \| IDinTask) |  |  |  |  |  |  |
| Intercept | -.095 | .04 | 229.00 | -2.61 | .01 | 2B |
| RespPhaseNumeric | -.016 | .01 | 1238.00 | -2.10 | .04 | 2B* |
| beforeRespIBI (s) | .10 | .04 | 227.00 | 2.28 | .02 | 2B* |
| **Model 3B:** SemDis ~ RespPhaseNumeric × beforeRespIBI + (1 \| Pnum) + (1 \| IDinTask) | | | | | | |
| Intercept | -.092 | .04 | 372.00 | -2.16 | .03 | 3B |
| RespPhaseNumeric | -.025 | .06 | 1247.00 | -0.41 | .69 | 3B |
| beforeRespIBI (s) | .10 | .05 | 374.00 | 1.87 | .06 | 3B |
| RespPhaseNumeric × beforeRespIBI | .01 | .08 | 1247.00 | .14 | .89 | 3B |
| **Model 1C:** SemDis ~ RespPhaseNumeric + (1\|Pnum) + (1\|IDinTask) |  |  |  |  |  |  |
| Intercept | .01 | .00 | 115.30 | 3.44 | .00 | 1C |
| RespPhaseNumeric | .01 | .00 | 1372.00 | 1.37 | .17 | 1C |
| **Model 2C:** SemDis ~ RespPhaseNumeric + beforeRespIBI + (1\|Pnum) + (1\|IDinTask) |  |  |  |  |  |  |
| Intercept | .02 | .02 | 205.70 | .91 | .37 | 2C |
| RespPhaseNumeric | .01 | .00 | 1368.00 | 1.35 | .18 | 2C |
| beforeRespIBI (s) | -.0069 | .02 | 204.70 | -.321 | .75 | 2C |

**Table 2. Table of mixed effects models examining the relationship between SemDis and Cardiac Phase.**
|  |  |  |  |  |  |  |
| --- | --- | --- | --- | --- | --- | --- |
| Intercept | .01 | .02 | 392.50 | .45 | .65 | 3C |
| RespPhaseNumeric | .02 | .03 | 1379.00 | .67 | .51 | 3C |
| beforeRespIBI (s) | .00 | .03 | 393.80 | .03 | .98 | 3C |
| RespPhaseNumeric × beforeRespIBI | -.017 | .04 | 1379.00 | -.495 | .62 | 3C |

In contrast, response time (i.e. the inter-response interval) showed consistent and condition-specific relationships with CD (see Table 3). For the AUT, adding a quadratic term to the linear mixed-effects model significantly improved model fit (Δχ² (1) = 6.04, *p* = .010), reducing AIC (from 22436 to 22432). Both the linear (β = 3.07, *p* < .001) and the quadratic components (β = .29, *p* = .010) were significant. These results suggest that increased CD is associated with longer response times, with a stronger effect emerging at higher levels of deceleration (see Fig. 2A). Interpreted in physiological units, for a 1ms increase in CD, there was a 30-35 ms increase in response time, but for a 20 ms increase in CD, there was a 600-700 ms increase in response time. The fixed effects explained a modest proportion of the variance (*R²_m* = .06), the full model with random effects (participant and trial number) performed better (*R²_c* = .19).

**Figure 2.**
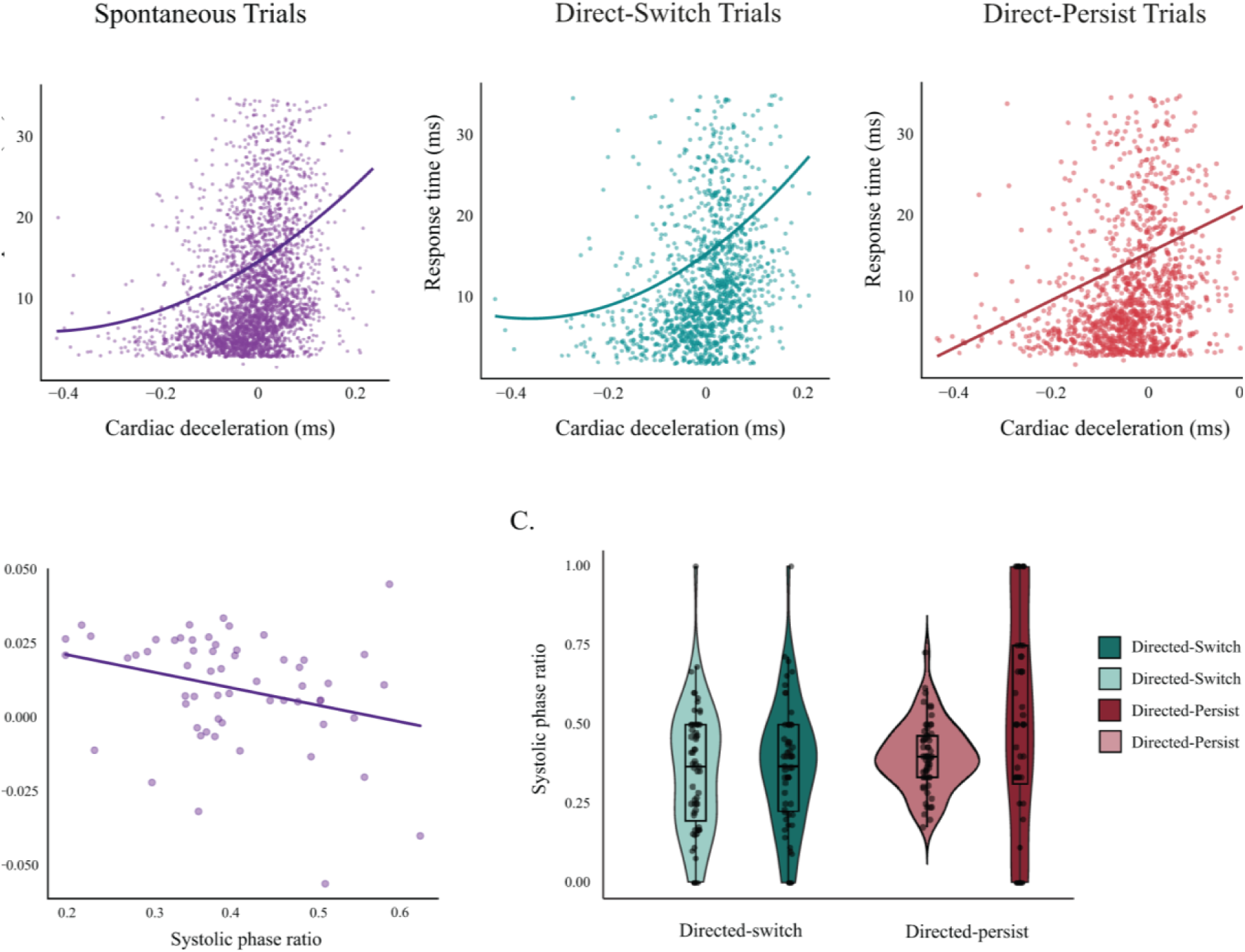
Condition dependent association between cardiac activity and ideation. **A)** Mixed-effects modelling of the relationship between cardiac deceleration and response time for all conditions. The left (purple) graph shows the quadratic relationship between cardiac deceleration and response time during spontaneous ideation. The middle (teal) graph shows the same during directed-switching trials. The graph on the right (red) shows the linear relationship during directed-persisting trials. **B)** The relationship between semantic distance and systolic phase ratios for the spontaneous condition. These are averaged by participant for the spontaneous condition. **C)** Differences in systolic phase ratios by directed instruction (switch/persist) and accuracy (hit/miss). The directed-switch conditions are again represented in teal, and the persist conditions are represented in red. The darker shading denotes the missed trials, while the lighter shading represents the accurate (hit) trials.

**Table 3.**
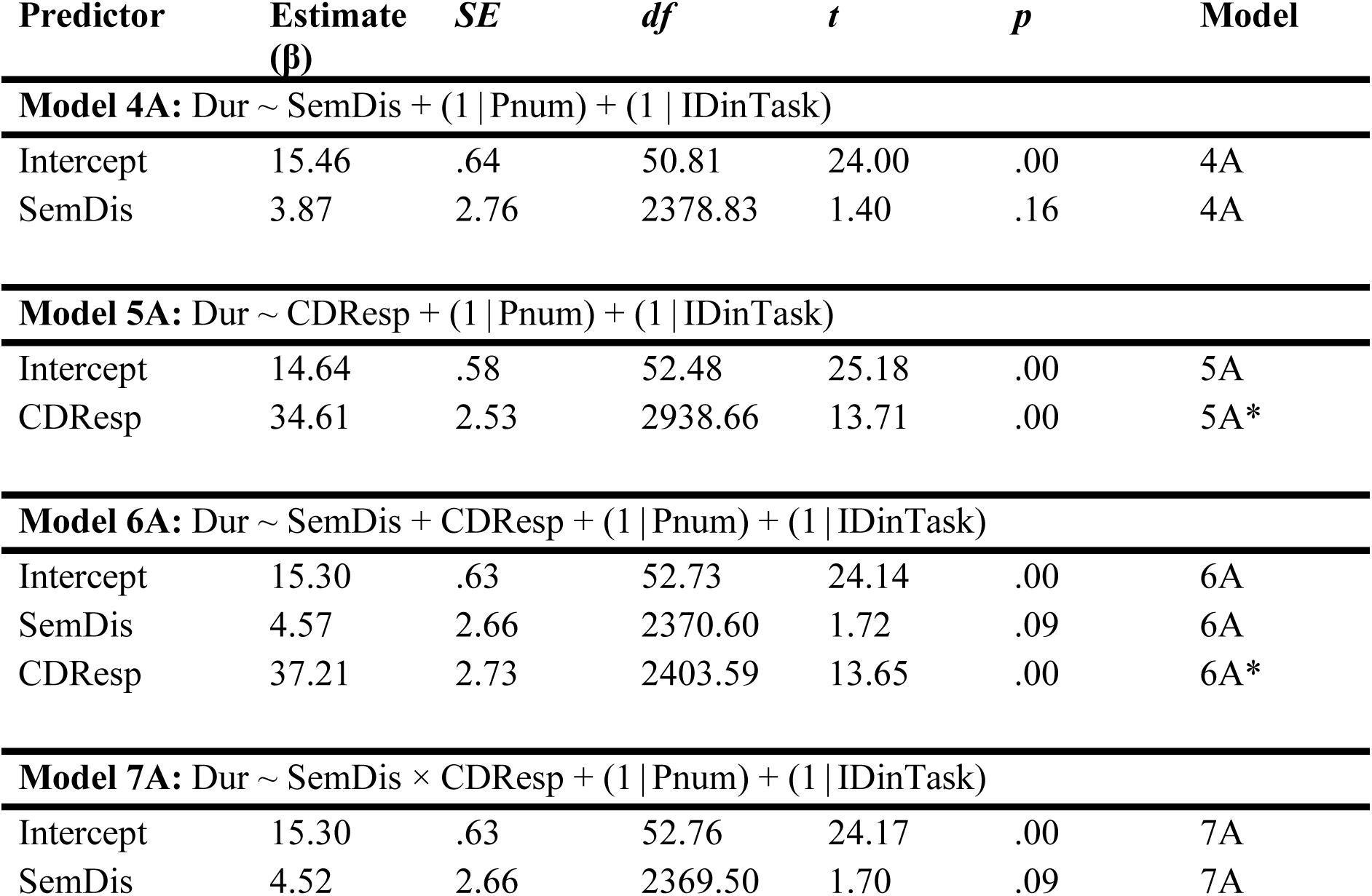
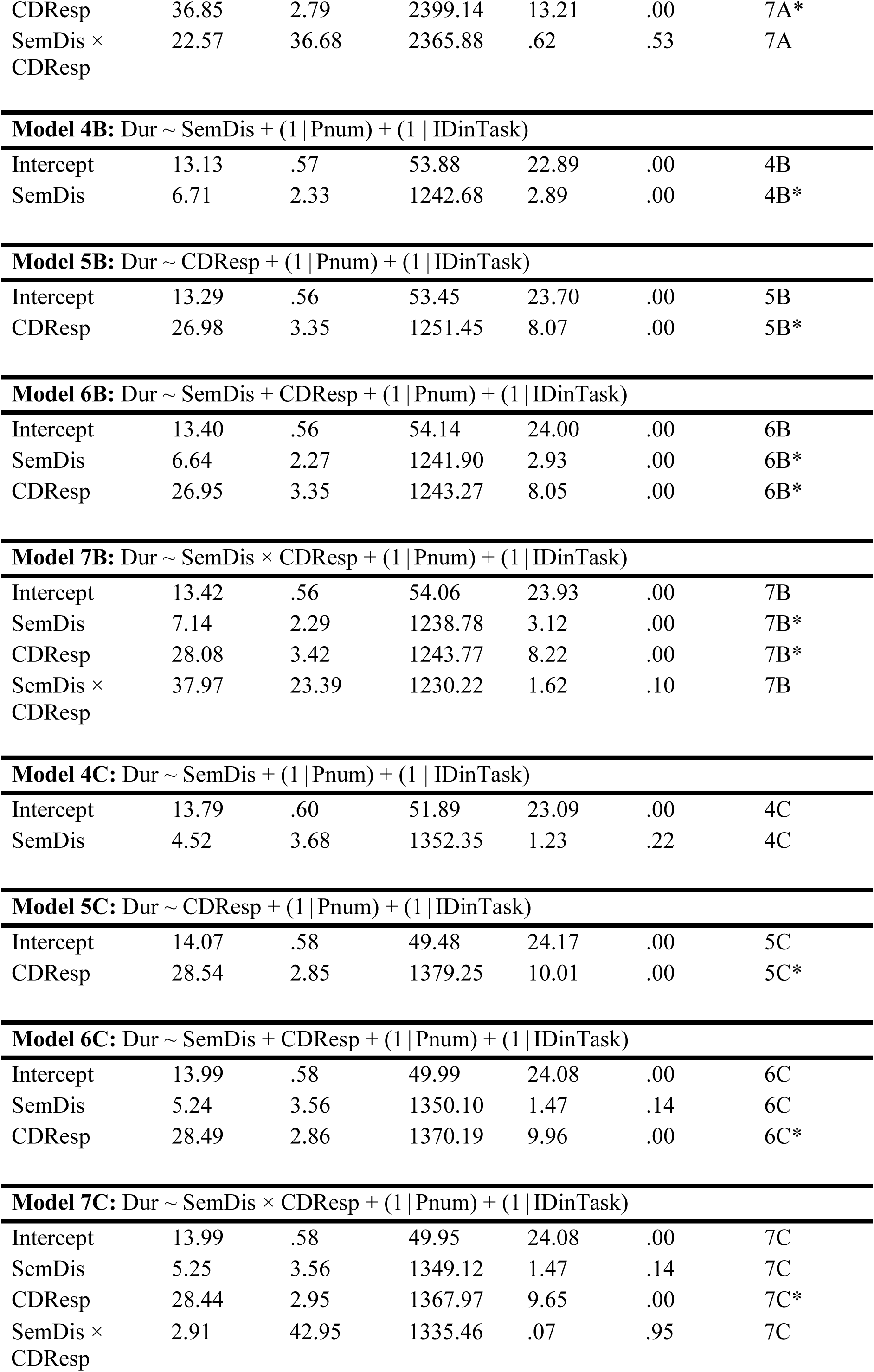
Mixed effects models examining the relationship between response time and Cardiac Deceleration. Model numbers reflect model structure (Model 4: SemDis; Model 5: CDResp; Model 6: SemDis + CDResp; Model 7: SemDis × CDResp), whereas letters denote experimental condition (A = spontaneous, B = directed persist, C = directed switch). All models include random intercepts for participants (Pnum) and trials (IDinTask). These linear mixed-effects models were run using two fixed factors, semantic distance (SemDis) and cardiac deceleration during response (CDResp), on the duration of the thinking period from the moment they ended the last trial to the moment they pressed the button to respond (Dur), using participant ID (Pnum) and response number (IDinTask) as random factors. They also test the potential interaction effects (×) between the fixed factors.

| Predictor | Estimate<br>($\beta$) | SE | df | t | p | Model |
| --- | --- | --- | --- | --- | --- | --- |
| **Model 4A:** $\text{Dur} \sim \text{SemDis} + (1\|\text{Pnum}) + (1\|\text{IDinTask})$ |  |  |  |  |  |  |
| Intercept | 15.46 | .64 | 50.81 | 24.00 | .00 | 4A |
| SemDis | 3.87 | 2.76 | 2378.83 | 1.40 | .16 | 4A |
| **Model 5A:** $\text{Dur} \sim \text{CDResp} + (1\|\text{Pnum}) + (1\|\text{IDinTask})$ |  |  |  |  |  |  |
| Intercept | 14.64 | .58 | 52.48 | 25.18 | .00 | 5A |
| CDResp | 34.61 | 2.53 | 2938.66 | 13.71 | .00 | 5A* |
| **Model 6A:** $\text{Dur} \sim \text{SemDis} + \text{CDResp} + (1\|\text{Pnum}) + (1\|\text{IDinTask})$ |  |  |  |  |  |  |
| Intercept | 15.30 | .63 | 52.73 | 24.14 | .00 | 6A |
| SemDis | 4.57 | 2.66 | 2370.60 | 1.72 | .09 | 6A |
| CDResp | 37.21 | 2.73 | 2403.59 | 13.65 | .00 | 6A* |
| **Model 7A:** $\text{Dur} \sim \text{SemDis} \times \text{CDResp} + (1\|\text{Pnum}) + (1\|\text{IDinTask})$ |  |  |  |  |  |  |
| Intercept | 15.30 | .63 | 52.76 | 24.17 | .00 | 7A |
| SemDis | 4.52 | 2.66 | 2369.50 | 1.70 | .09 | 7A |
| CDResp | 36.85 | 2.79 | 2399.14 | 13.21 | .00 | 7A* |
| SemDis ×<br>CDResp | 22.57 | 36.68 | 2365.88 | .62 | .53 | 7A |
| **Model 4B:** Dur ~ SemDis + (1 \| Pnum) + (1 \| IDinTask) |  |  |  |  |  |  |
| Intercept | 13.13 | .57 | 53.88 | 22.89 | .00 | 4B |
| SemDis | 6.71 | 2.33 | 1242.68 | 2.89 | .00 | 4B* |
| **Model 5B:** Dur ~ CDResp + (1 \| Pnum) + (1 \| IDinTask) |  |  |  |  |  |  |
| Intercept | 13.29 | .56 | 53.45 | 23.70 | .00 | 5B |
| CDResp | 26.98 | 3.35 | 1251.45 | 8.07 | .00 | 5B* |
| **Model 6B:** Dur ~ SemDis + CDResp + (1 \| Pnum) + (1 \| IDinTask) |  |  |  |  |  |  |
| Intercept | 13.40 | .56 | 54.14 | 24.00 | .00 | 6B |
| SemDis | 6.64 | 2.27 | 1241.90 | 2.93 | .00 | 6B* |
| CDResp | 26.95 | 3.35 | 1243.27 | 8.05 | .00 | 6B* |
| **Model 7B:** Dur ~ SemDis × CDResp + (1 \| Pnum) + (1 \| IDinTask) |  |  |  |  |  |  |
| Intercept | 13.42 | .56 | 54.06 | 23.93 | .00 | 7B |
| SemDis | 7.14 | 2.29 | 1238.78 | 3.12 | .00 | 7B* |
| CDResp | 28.08 | 3.42 | 1243.77 | 8.22 | .00 | 7B* |
| SemDis ×<br>CDResp | 37.97 | 23.39 | 1230.22 | 1.62 | .10 | 7B |
| **Model 4C:** Dur ~ SemDis + (1 \| Pnum) + (1 \| IDinTask) |  |  |  |  |  |  |
| Intercept | 13.79 | .60 | 51.89 | 23.09 | .00 | 4C |
| SemDis | 4.52 | 3.68 | 1352.35 | 1.23 | .22 | 4C |
| **Model 5C:** Dur ~ CDResp + (1 \| Pnum) + (1 \| IDinTask) |  |  |  |  |  |  |
| Intercept | 14.07 | .58 | 49.48 | 24.17 | .00 | 5C |
| CDResp | 28.54 | 2.85 | 1379.25 | 10.01 | .00 | 5C* |
| **Model 6C:** Dur ~ SemDis + CDResp + (1 \| Pnum) + (1 \| IDinTask) |  |  |  |  |  |  |
| Intercept | 13.99 | .58 | 49.99 | 24.08 | .00 | 6C |
| SemDis | 5.24 | 3.56 | 1350.10 | 1.47 | .14 | 6C |
| CDResp | 28.49 | 2.86 | 1370.19 | 9.96 | .00 | 6C* |

**Table 3. Mixed effects models examining the relationship between response time and Cardiac Deceleration.**
|  |  |  |  |  |  |  |
| --- | --- | --- | --- | --- | --- | --- |
| Intercept | 13.99 | .58 | 49.95 | 24.08 | .00 | 7C |
| SemDis | 5.25 | 3.56 | 1349.12 | 1.47 | .14 | 7C |
| CDResp | 28.44 | 2.95 | 1367.97 | 9.65 | .00 | 7C* |
| SemDis ×<br>CDResp | 2.91 | 42.95 | 1335.46 | .07 | .95 | 7C |

For the directed-persist condition, adding a quadratic term did not improve model fit (Δχ² (1) = 1.07, *p* = .300), nor was there a reduction in AIC (from 9513 to 9514). Still, the linear effect of CD remained significant (β = 26.98, *p* < .001), suggesting that even under directed persistence, CD predicted prolonged response times (Fig 2A). On the other hand, for the directed-switch condition, adding a quadratic term significantly improved model fit, (Δχ² (1) = 4.93, *p* = .030), and reduced AIC (from 10,183 to 10,180). Both the linear (β = 2.81, *p* < .001) and the quadratic term (β = .32, *p* = .030) were significant, indicating a similar nonlinear relationship between CD and response time as observed in the spontaneous condition. Notably, this model explained more variance (R*²_m* = .07; *R*²_*c* = .21), suggesting more stable individual-level associations in this condition (Fig. 2A).

Taken together, these findings partially supported our first hypothesis, indicating that greater CD robustly predicted response times across spontaneous, directed-persist, and directed-switch conditions, especially at higher levels of CD. However, CD did not consistently predict inter-response semantic distance, suggesting that cardiac-behavior interactions may be more closely related to the temporal dynamics of ideational search than to semantic remoteness.

### Individual differences in cardiac phase bias and ideation behaviors

To examine whether individual differences in cardiac phase ratio (responses given during systole vs. diastole) was associated with inter-response semantic distance, we first focused on the AUT. A linear regression revealed that the ratio of systolic phase timing during ideation was a significant predictor of average semantic distance, where systolic biases were associated with lower semantic distance (β = −.06, *t* (57) = −2.33, *p* = .020). This model explained 9% of the variance in semantic distance (*R²* = .09; see Fig. 2B).

To evaluate whether the relationship between cardiac phase ratios and inter-response semantic distance was moderated by other factors, we conducted multiple regressions incorporating average heart rate and average response time as potential moderators. Neither variable significantly interacted with cardiac phase ratio in predicting semantic distance (*ps* > .100), indicating that the relationship does not vary according to baseline heart rate or behavioral response time.

We next examined whether similar associations between cardiac phase ratio and inter-response semantic distance were present in the D-AUT. No significant relationship emerged in the directed-persist condition (β = -.05, *t* (57) = -1.39, *p* = .190) or the directed-switch condition (β = .02, *t* (57) = .69, *p* = .490), indicating that phase ratio effects were attenuated or absent when participants followed externally imposed strategies. However, when controlling average IBI length, a proxy for tonic heart rate, we found that in the directed-persist condition, heart rate was significantly associated with semantic distance, β = .13, *t* (57) = 2.41, *p* = .020, such that slower heart rates were linked to more semantically distant responses. This suggests that while cardiac phase ratio is associated with spontaneous AUT ideation, broader cardiac state variables like heart rate may relate to performance under constrained cognitive contexts.

To further probe how cardiac phase ratio interacted with D-AUT task instructions and behavioral performance, a two-way ANOVA was conducted with condition (directed-switch vs. directed-persist) and task accuracy (correct vs. incorrect responses: hit vs. miss) as factors, predicting the proportion of systolic responses (see Fig. 2C). The main effect of condition was significant (*F* (1, 225) = 9.15, *p* = .003), with the directed-switch condition (*M* = .45, *SD* = .28) eliciting a higher proportion of systolic responses than the directed-persist condition (*M* = .36, *SD* = .20). The main effect of task accuracy was not significant (*F* (1, 225) = 2.09, *p* = .150), indicating that cardiac phase ratio did not significantly vary across correct and incorrect trials. However, the interaction between condition and task accuracy was marginally significant (*F* (1, 225) = 4.35, *p* = .038). Follow-up pairwise comparisons revealed that when participants were instructed to persist in the same category (directed-persist), they responded during systole 35% of the time when responding correctly (persist_hit; *M* = .35, *SD* = .20), and 45% of the time when responding incorrectly (persist_miss; *M* = .45, *SD* = .20). Likewise, when participants were instructed to switch categories (directed-switch), they responded during systole 40% of the time when responding correctly (switch_hit; *M* = .40, *SD* = .11) and 51% of the time when responding incorrectly (switch_miss; *M* = .51, *SD* = .34). Taken as a whole, these findings suggest that higher systolic dominance was associated with a reduced ability to follow switch or persist instructions during the D-AUT.

To control for potential confounds related to response time, we also conducted a two-way ANOVA with condition (directed-switch vs. directed-persist) and task accuracy (correct vs. incorrect responses: hit vs. miss) as factors predicting response time. No significant main effects or interactions emerged (*ps* ≥ .050), indicating no significant differences in response time by condition or accuracy.

Altogether, these analyses provided partial support for our second hypothesis; that ideational transitions would be timed to specific cardiac phases. During the AUT, greater inter-response semantic distance was associated with greater diastolic dominance while shorter inter-response semantic distance was associated with greater systolic dominance. These behavioral analyses indicate that associations between cardiac phase and ideation are apparent at the level of condition-averaged tendencies, rather than as a reliable trial-level predictor of semantic distance or response time. Thus, the behavioral findings are interpreted as reflecting stable individual differences and condition-dependent biases in cardiac timing. The neurophysiological analyses that follow are presented as complementary evidence aimed at characterizing potential neural mechanisms that may underlie these behavioral patterns.

### Time-frequency correlates of cardiac timing during spontaneous ideation

To examine the relationship between cardiac timing and neural activity during spontaneous idea generation, we conducted linear regressions linking cardiac measures (cardiac deceleration and cardiac phase ratio) to EEG amplitude changes to the parametric SemDis regressor (inter-response semantic distance) for the AUT condition obtained in our convolution GLM analysis. The mean time-frequency amplitudes were extracted from three frequency bands: theta (4 – 6 Hz), alpha (8 – 12 Hz), and beta (13 – 30 Hz); averaging over a pre-event window of interest (-2 to 0 s) for each regressor at nine bilateral regions of interest (see *Methods* section).

CD was significantly associated with alpha desynchronization in posterior-occipital (β = -102233.23, *p_*FDR = .030) and occipital (β = -97756.82, *p_*FDR = .030) regions during the AUT (Fig. 3A). Exploratory analyses using originality scores (originality is assessed as the semantic distance from the cue word to the response) – instead of inter-response semantic distance scores – as convolution GLM regressors, revealed similar associations with alpha desynchronization in posterior-occipital (β = -109060.68, *p* < .001) and occipital (β = - 116676.00, *p* = .008) regions. In addition, alpha synchronization in centro-temporal regions was positively associated with CD (β = 43307.76, *p* = .008). Together, these findings indicate that stronger CD was specifically associated with enhanced alpha suppression in posterior visual processing regions during spontaneous ideation (see Fig. 3B; also Supplementary Figure 1).

**Figure 3.**
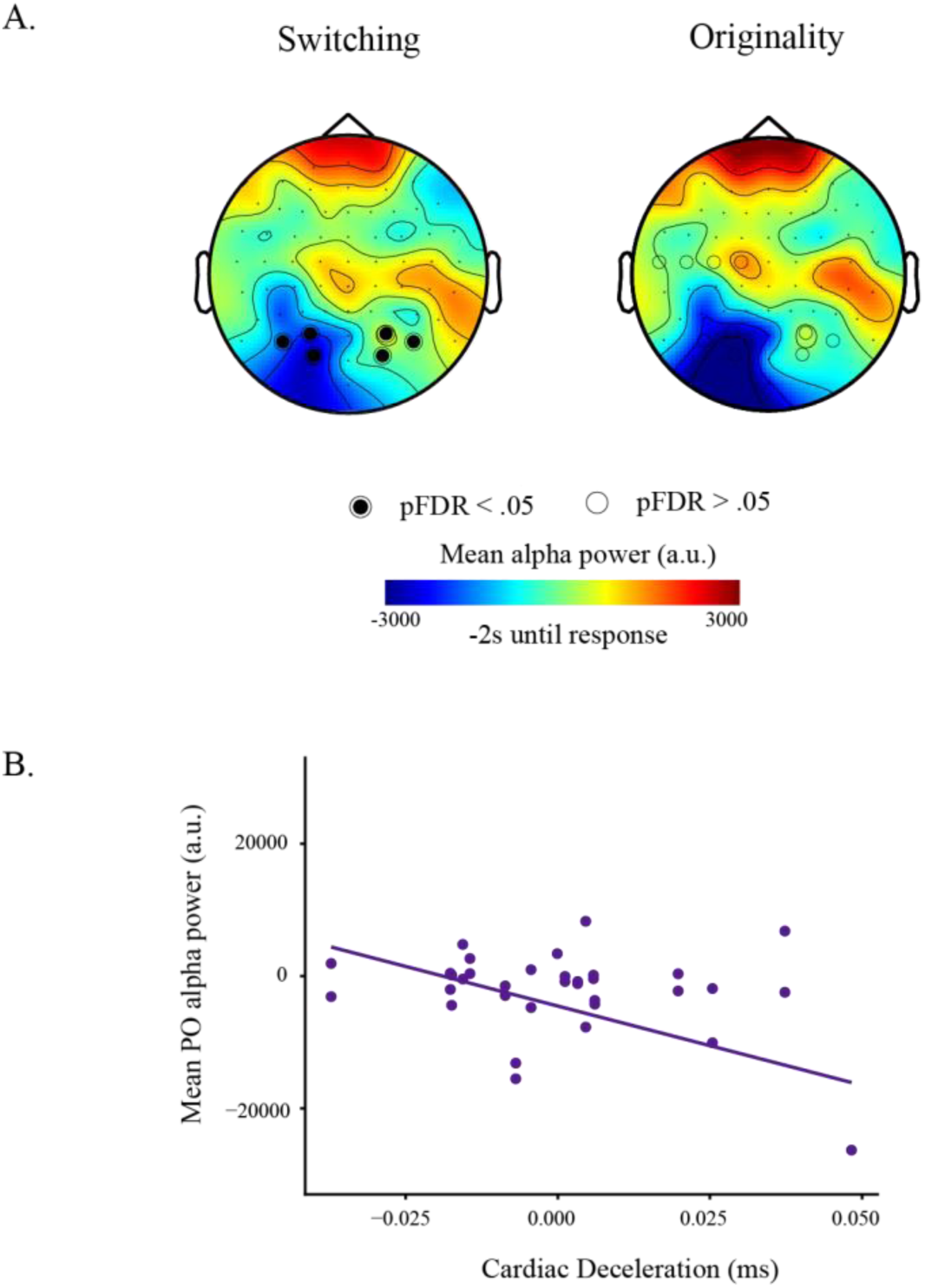
Cardiac deceleration linked to posterior-occipital alpha desynchronization during spontaneous ideation. **A)** The significant relationship between cardiac deceleration (ms) and average parieto-occipital alpha power in arbitrary units (a.u.) during spontaneous (AUT) ideation. **B)** Topology of averaged alpha power during spontaneous (AUT) ideation. The left topoplot shows alpha power from the convolution GLM using inter-response semantic distance (switching) as a parametric regressor, while the right topoplot shows alpha power from the convolution GLM using cue-response semantic distance (originality) as a parametric regressor. Topology was derived from -2 seconds until response. Greater cardiac deceleration was associated with lower parieto-occipital alpha power. Highlighted electrodes represent ROIs showing a significant relationship (*p* < .05 uncorrected is highlighted with a simple circle; *pFDR* < .05 corrected is highlighted with the circle filled in) with cardiac deceleration.

Further exploratory analyses with the GLM using originality scores as a parametric regressor instead of inter-response semantic distance found that beta synchronization in anterio-frontal (β = 12398.57, *p* = .010) and parieto-occipital (β = 17547.00, *p* = .010) regions were associated with the tendency to respond during systole in the spontaneous condition. There was no significant relationship between cardiac phase ratio or CD and beta band activity linked to inter-response semantic distances. This pattern suggests that systolic response bias co-occurred with elevated beta activity in frontal-parietal regions when participants generated more original ideas, rather than when they explored or exploited the previous idea.

When testing if theta activity linked to spontaneous inter-response semantic distance was associated with CD, we found no significant relationship. Interestingly, we found that theta synchronization in anterior mid-frontal regions linked to spontaneous inter-response semantic distance (β = 45308.71, *p* < .001, *p*_FDR = .002) was significantly related to the tendency to respond during systole (cardiac phase ratio; see Fig. 4). Exploratory analyses revealed that this pattern was also reflected in originality (β = 34839.77, *p* = .002). These effects are consistent with anterior mid-frontal involvement in monitoring, metacognitive appraisal, or internally directed attention.

**Figure 4.**
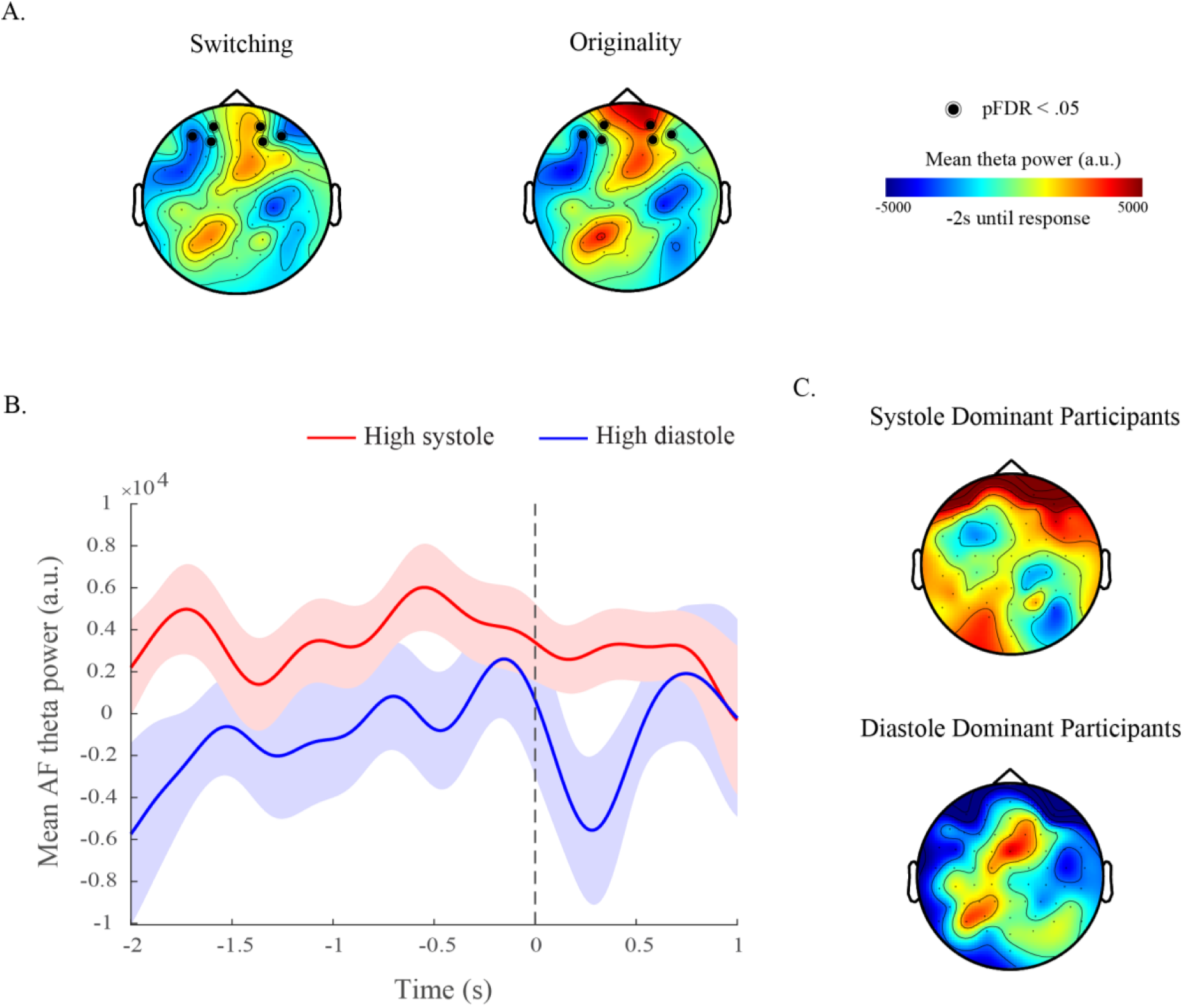
Cardiac phase linked to theta power during spontaneous condition. **A)** Topology of averaged theta power during spontaneous (AUT) ideation. The left topoplot shows inter-response semantic distance (switching), while the right topoplot shows the cue-response semantic distance (originality). Highlighted electrodes denote ROIs which were significantly (pFDR < .05) associated with cardiac phase ratios. Topology was derived from -2 seconds until response (0). Greater frontal theta was associated with the tendency to respond during systole. **B)** Averaged timeseries of frontal theta power 2 seconds before and one second after the response (0) for high systole (red) and high diastole (blue) participants during spontaneous (AUT) ideation, defined as the top 20 highest systole ratio and the lowest systole ratio. The high-systole group (M = 2046.03, SD = 8187.07, n = 18) showed significantly greater mean frontal theta power than the high-diastole group (M = -4567.05, SD = 9058.97, n = 18) across the analysis window of -2 seconds to response (0), *t*(34) = 2.30, *p* = .028, d = 0.77. The shaded regions represent +/- 1 SEM. **C)** Averaged theta power topology for high systole (top) and high diastole (low) participants during spontaneous ideation. Highlighted electrodes denote ROIs which were significantly (pFDR < .05) associated with cardiac phase ratios. Topology was derived from -2 seconds until response (0).

To ensure that the relationship between cardiac timing and ideation-related neural activity was not driven by movement-related neural activity, we performed supplementary analyses including the EEG activity obtained from the second regressor of our GLM model for the AUT condition: button-press (see Supplementary Figure 2). We conducted these analyses to account for the *general inhibition* hypothesis posed by Obrist (1968), which proposed that CD reflects reduced peripheral muscular activity during action execution rather than attentional modulation. Results indicated that beta-band suppression during button-presses was robustly associated with cardiac acceleration across multiple regions (see *Supplementary Analysis*). Together, these analyses revealed dissociable patterns: motor-related activity was characterized by beta-band modulation and cardiac acceleration consistent with parasympathetic withdrawal during action execution (Skora et al., 2022; Hashemi et al., 2019), whereas ideation-related activity showed alpha desynchronization coupled with cardiac deceleration. These distinct profiles indicate that the cardiac-heart rate changes coupled to EEG time-frequency activity observed in ideation analyses cannot be fully attributed to motor confounds.

Altogether, the data provided partial support for our third hypothesis. For the AUT, results show that cardiac–behavioral interactions were mirrored in frequency-domain activity in the alpha and theta bands. Greater inter-response semantic distance was linked to posterior-occipital alpha desynchronization and mid-frontal theta synchronization. The former activity was coupled to higher CD, as expected. However, the latter was linked to responses timed to systole, when the opposite was expected. Additionally, exploratory analyses on beta band activity revealed that the amplitudes linked to inter-response semantic distance (and originality) are distinct from motor-related brain activity. Further exploratory analyses revealed neural-cardiac activities differentially linked to inter-response semantic distance and originality.

### Time-frequency correlates of cardiac timing during directed ideations

To examine the relationship between cardiac timing and neural activity during directed conditions, we applied the same methods as before. In contrast to the spontaneous condition, the changes in time-frequency amplitude were obtained using convolution GLM with four discrete regressors (switch_hit, switch_miss, persist_hit, persist_miss) for the D-AUT.

For the alpha band, only one significant association passed FDR correction: in correct directed-switch (switch_hit) trials, greater alpha synchronization in anterio-frontal regions was positively associated with the tendency to respond during systole (β = 14.38, *p_*FDR = .040, see Fig. 5). This suggests that during accurately executed switching trials, systolic response bias co-occurred with increased frontal alpha activity. No other significant associations (FDR corrected) with cardiac timing were observed for alpha activity in the other three conditions (switch_miss, persist_hit, persist_miss). There was no significant relationship between cardiac timing and activity in the theta band for any of the directed conditions (all *ps* > .100).

**Figure 5.**
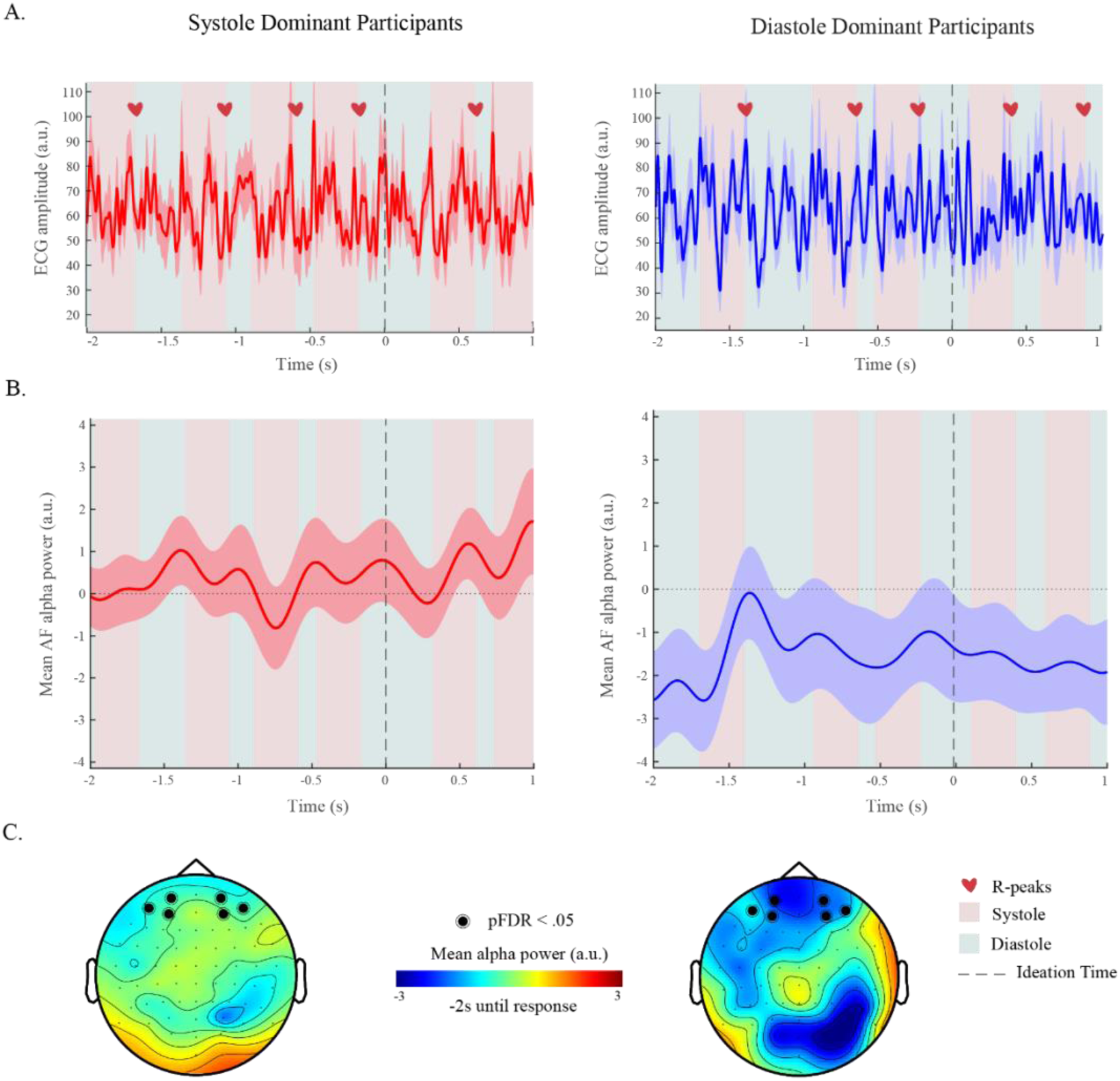
Cardiac phases linked to alpha activation during accurate directed switching. **A)** Event-locked cardiac time series across the accurate directed-switching trials. Displayed are grand-average ECG waveforms time-locked to response onset (0 s – dotted line), plotted separately for high-diastole (blue/right) and high-systole (red/left) participants for visualization, defined as the top 20 highest systole ratio and the lowest systole ratio. Shaded error bands denote the standard error of the mean. The heart icons denote approximate R-peak locations identified from local maxima of the averaged trace. Background shading indicates canonical cardiac phases derived from approximate R-peak locations (denoted by a heart icon) in the averaged waveform, with systole defined as the 300 ms interval following each R-peak and diastole defined as the interval from the end of systole to the subsequent R-peak. ECG power is shown in arbitrary units (a.u.), and the time axis spans from 2 s before to 1 s after the response. **B)** Averaged timeseries of parieto-occipital alpha power for high-systole (red/left) and high-diastole (blue/right) participants during accurate directed-switching (D-AUT). The shaded regions represent +/-1 SEM. An independent-samples *t*-test revealed a significant difference between groups, *t*(34) = −2.63, *p* = .013, Cohen’s d = −0.88. Participants in the high diastole group (M = −5607.45, SD = 13241.87, n = 18) exhibited significantly lower AF alpha power compared to those in the high systole group (M = 4572.94, SD = 9726.72, n = 18). **C)** Averaged alpha power topology for high-systole (left) and high-diastole (right) participants during accurate directed-switching. Highlighted electrodes represent ROIs with a significant (pFDR < .050) relationship between alpha power and cardiac phase ratio.

In the beta band, exploratory analyses revealed a weak association between systolic response tendency and beta synchronization in accurate directed-persist (persist_hit) trials: in parieto-temporal (β = 3.14, *p* = .020) and parieto-occipital (β = 3.57, *p* = .020) regions.

For the D-AUT, the data partially supported our third hypothesis: that cardiac–behavioral interactions would be mirrored in time-frequency activity in the alpha band. However, we did not find a similar effect in the theta band, as expected. Again, our results differed from our expectations. Specifically, the tendency to respond during systole – not diastole – was associated with alpha synchronization in anterio-frontal regions. This was only the case for correct directed-switching, and not for incorrect directed-switching or either correct or incorrect directed-persisting. Moreover, CD was not significantly associated with oscillatory activity in any of the frequency bands for the directed condition, which was another unexpected result.

### Circular analysis of mean cardiac response phase and ideation outcomes

Given the inherently cyclical nature of cardiac activity, we also employed the widespread method of circular statistics to assess the timing of responses across the cardiac cycle (see *Methods* section). The duration of systolic and diastolic phases are not equivalent, and may change based on heart rate (Fridericia, 2003). To examine whether participants exhibited non-random temporal alignment of responses within the cardiac cycle, we conducted Rayleigh tests of uniformity across ideation conditions. As summarized in Table 4, mean response phases during spontaneous, directed-persist, and directed-switch conditions did not significantly deviate from a uniform distribution (all *ps* > .100), indicating no consistent group-level phase-locking across individuals.

**Table 4.**
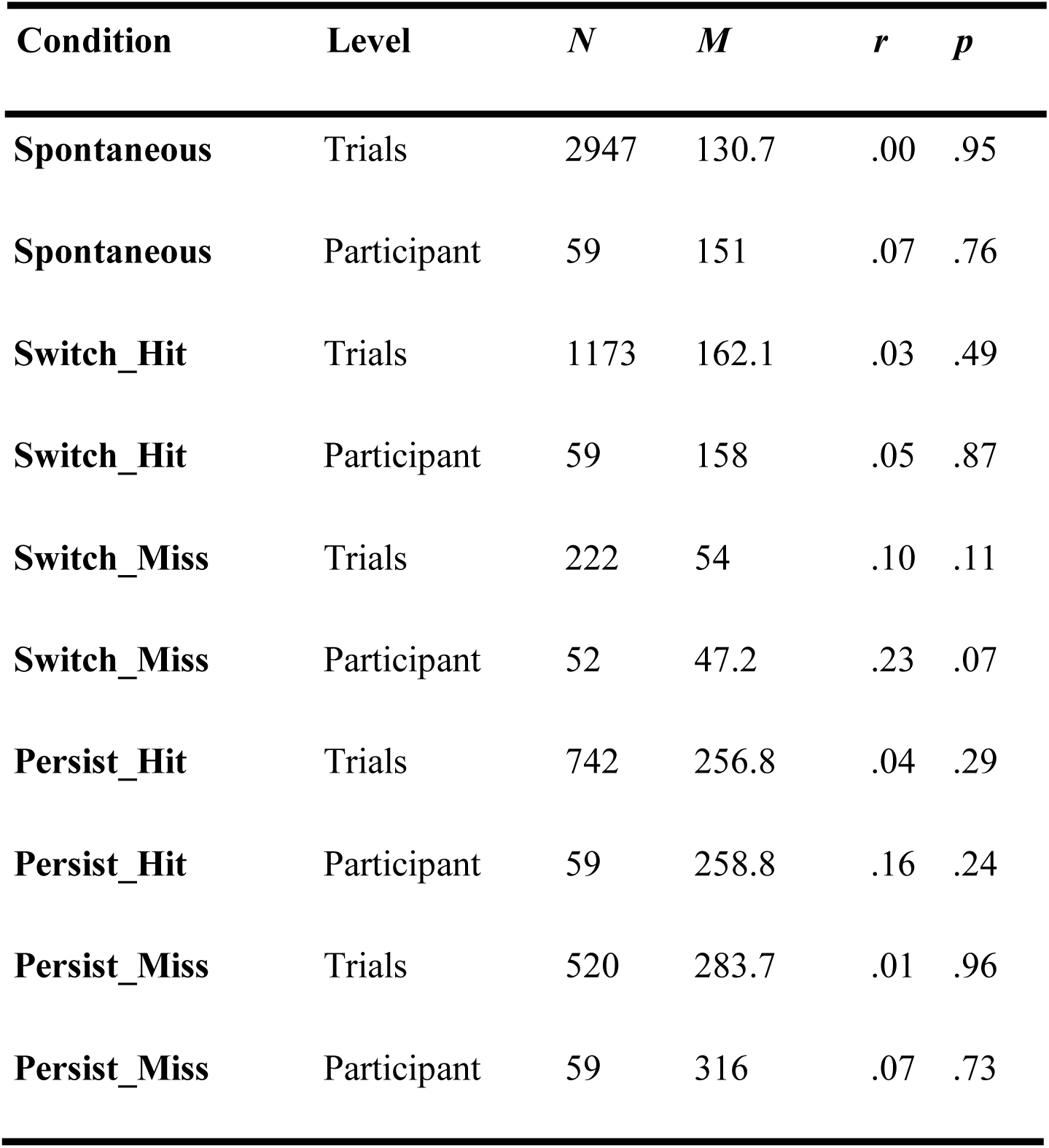
Raleigh tests for vector length and position in the cardiac cycle of response for all conditions at the trial and participant levels. Trial- and participant-level Raleigh tests of mean degree of phase angle for each condition.

We next explored whether individual differences in mean phase angle (i.e., average position within the cardiac cycle during button-press expressed in radians) and, separately, phase concentration (i.e., how loosely or tightly phase-locked responses were to a specific point in their cardiac cycle; irrespectively of phase) related semantic distance. Circular-linear correlation analyses revealed no significant associations in any of the three ideation conditions (all *ps*_FDR > .100), meaning that there was no significant relationship between position in the cardiac phase during ideation and the inter-response semantic distance.

When considering inter-individual variability in phase concentration, we observed only one exception. In correct directed-persist (persist_hit) trials, participants with responses more tightly clustered within their cardiac cycle exhibited greater inter-response semantic distances (*r* = .354, 95% CI [.108, .560], p < .001). However, this association was specific to the directed-persist correct condition and did not generalize across conditions after FDR correction.

### Time-frequency correlates of cardiac phase concentrations

To further account for the cyclical nature of the cardiac cycle regarding EEG amplitudes, we investigated whether individual differences in phase concentration was associated with neural activity during creative ideation. In other words, if the clustering of response timing within the cardiac cycle was associated with EEG time-frequency during ideation for all conditions.

Significant positive correlations emerged for correct directed-persist trials. Across multiple regions of interest (ROIs), greater alpha synchronization was significantly associated with higher phase concentration, indicating that individuals whose responses clustered more tightly within the cardiac cycle (irrespective of phase) also exhibited enhanced alpha activity. The effect was observed across regions (see Fig. 6): occipital (*r* = .50, *p* < .001, *p_*FDR = .004), anterio-frontal (*r* = .48, *p* < .001, *p_*FDR = .004), parieto-occipital (*r* = .48, *p* < .001, *p_*FDR = .004), frontal (*r* = .43, *p* = .002, *p_*FDR = .018), and parieto–temporal (*r* = .42, *p* = .002, *p_*FDR = .018). These effects indicate a tight coupling between the timing regularity of responses within cardiac cycles and elevated alpha activity during successful semantic exploitation.

**Figure 6.**
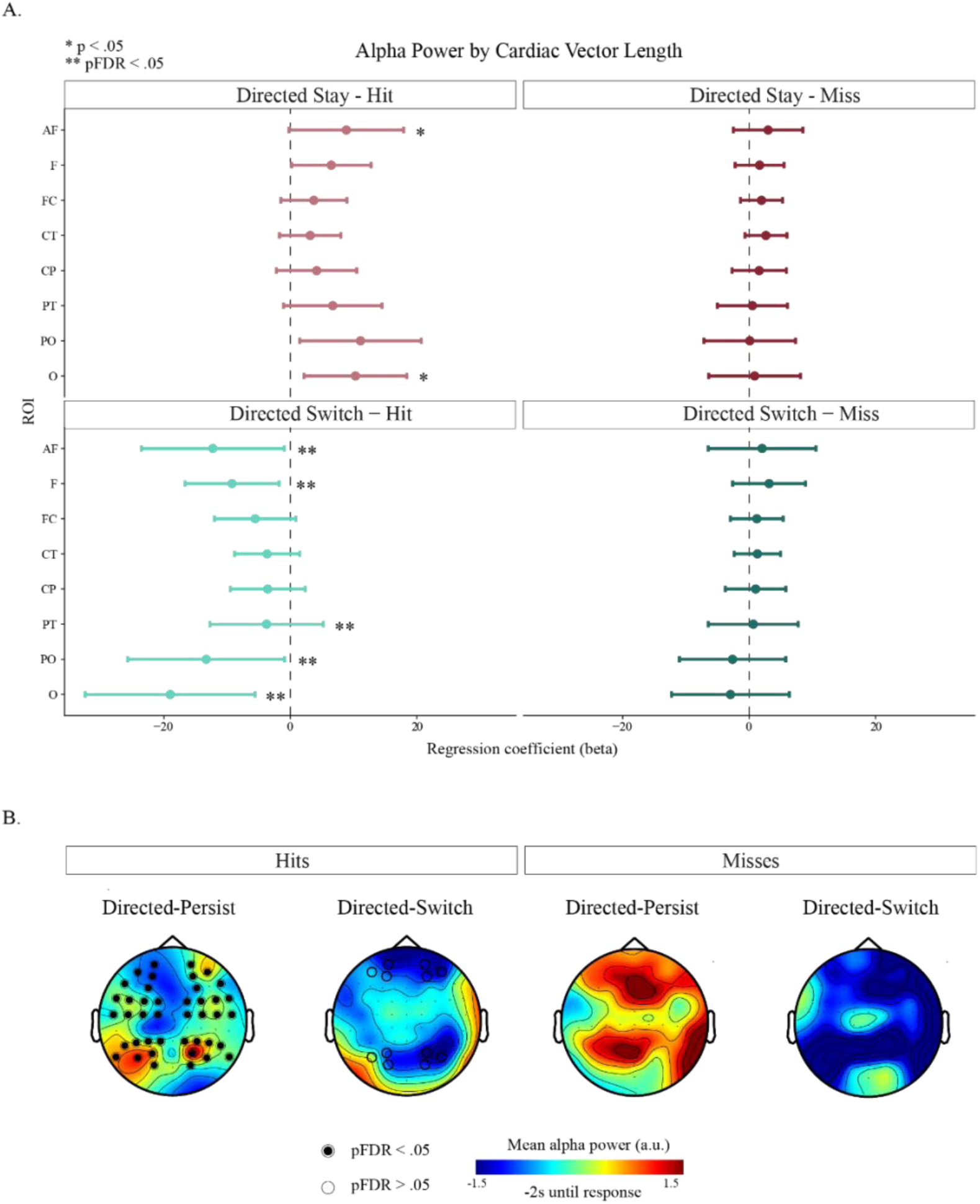
Cardiac *phases linked to alpha activation during accurate directed conditions*. **A)** Regression coefficients (β) from linear models predicting alpha-band power from cardiac phase concentration (vector length r) are shown for each region of interest (ROI) and task condition. Separate models were estimated for each ROI and directed (D-AUT) condition (persist_hit, persist_miss, switch_hit, switch_miss). Points indicate the estimated regression coefficient relating vector length to alpha power, and horizontal bars represent 95% confidence intervals. The dashed vertical line denotes zero effect. Positive coefficients indicate that greater cardiac phase concentration (i.e., responses occurring more consistently at a particular position within the cardiac cycle) is associated with increased alpha power within that ROI, whereas negative coefficients indicate the opposite relationship. Asterisks denote statistical significance (*p < .05 uncorrected; **FDR-corrected p < .05 across ROIs within each condition). **B)** Averaged alpha power topology for directed condition (from left to right: persist_hit/switch_hit and persist_miss/switch_miss). Highlighted electrodes represent ROIs showing a significant relationship (p < .05 uncorrected is highlighted with a simple circle; pFDR < .05 corrected is highlighted with the circle filled in) between alpha power and cardiac phase concentration (vector length r).

For correct directed-switch (switch_hit) trials, alpha amplitudes were negatively correlated with phase concentration in anterio-frontal (*r* = -.44, *p* = .001, *p_*FDR= .050) and occipital regions (*r* = -.37, *p* = .007, *p_*FDR = .160) regions, although these did not pass FDR correction. Thus, evidence for an inverse relationship during correct directed switching was limited and region-specific.

During spontaneous ideation, no associations between phase concentration and EEG band amplitudes passed FDR correction. Analysis indicated modest negative correlations between phase concentration and occipital alpha (*r* = -.32, *p* = .020) and theta (*r* = -.32, *p* = .020), and parieto-occipital beta (*r* = -.31, *p* = .020); however, these effects did not survive correction.

Together, these findings suggest a condition-specific relationship between interoceptive timing concentration and alpha-band activity. Greater phase concentration was associated with elevated alpha power during successful semantic exploitation, whereas associations in other contexts were weaker or did not survive strict statistical threshold. The failure to replicate previous analyses regarding cardiac phase timing encourages caution when interpreting the results.

### Cardiac phase and alpha synchrony

To further assess whether the timing of responses within the cardiac cycle was associated with time-frequency EEG activity, we examined the coupling between response phase angle (average position within the cardiac cycle during button-press expressed in radians) and EEG power. A significant circular-linear correlation emerged in the spontaneous ideation condition: the closer the response was to diastole, the greater the frontal alpha synchrony during switching trials (*r(cl)* = .45, χ²(2) = 10.61, *p* = .005). This finding provides support for our second hypothesis and third hypothesis, that frontal alpha activity associated with spontaneous exploration would be more likely to occur diastole.

A weaker association was observed in accurate directed-switch trials, where parieto-occipital alpha was modestly related to response phase (*r(cl)* = .34, χ²(2) = 6.15, *p* = .046), although this effect did not survive FDR correction (*p_*FDR = .610). No other significant phase-amplitude relationships were significant across ideation conditions or frequency bands after statistical correction.

## Discussion

This study provides the first systematic evidence that creative ideation is temporally aligned with rhythmic cardiac-brain coordination. By simultaneously recording cardiac and neural signals during spontaneous and directed creative ideation, we reveal four key findings that together establish embodied interoceptive effects on creative cognition: (i) cardiac deceleration (CD) robustly predicts response time in a condition-specific manner – nonlinear during spontaneous and directed switching, but linear during directed persisting – yet does not predict the semantic distance between responses, indicating that heart-rate slowing indexes cognitive effort rather than ideational content; (ii) cardiac phase ratios (i.e., the relative proportion of responses during systole versus diastole) are robustly associated with ideation behavior, with responses clustering toward diastolic phases during spontaneous semantic exploration and correct directed exploration, and toward systolic phases during spontaneous exploitation and directed exploitation; (iii) individual differences in cardiac phase ratios predicted mid-frontal theta synchronization during spontaneous semantic exploration, while, separately, successful directed exploitation co-occurred with widespread alpha synchronization and greater cardiac phase concentration; and (iv) the magnitude of CD is coupled with parieto-occipital alpha desynchronization during spontaneous semantic exploration.

Together, these findings offer an integrative account of how interoceptive rhythms interact with neural activity during creative ideational search in semantic space, expanding current cortico-centric models of creativity by positioning the heart-brain axis as an active contributor to divergent thinking. More broadly, they advance growing evidence that visceral rhythms continuously bias higher-order cognitive processes, positioning the body not merely as a substrate for cognition but as an active contributor to thought (e.g., Allen et al., 2019; Buzsáki, 2025; Park & Tallon-Baudry, 2014; Singer & Damasio, 2025).

### Cardiac deceleration as a marker of ideational effort, not content

We initially hypothesized that CD, a parasympathetic marker of adaptive resource allocation (Hashemi et al., 2019; Van Roon et al., 2004), would predict both spontaneous semantic exploration and accurate implementation of cued ideation behavior (switching vs. persistence), based on evidence that semantic switching is cognitively more demanding than persistence (Mastria et al., 2021; Mazza et al., 2023; Roelofs et al., 2010). We also predicted that CD would scale with response times, since longer response times during divergent thinking are often associated with greater effort and originality (Mazza et al., 2023).

Our results partially supported these predictions. CD robustly predicted response time across all conditions (β = 27–37 ms per 1 ms increase in inter-beat interval), with this association best captured by quadratic functions during exploration (both spontaneous and directed-switch explorations), but a linear relationship during directed-persistence. Notably, this relationship remained significant even when controlling for individual differences, serial-order effects, inter-response semantic distance, and the interaction between semantic distance and response time. The difference between semantic switching and persistence reveals condition-specific effort dynamics: exploration imposes escalating computational demands as individuals venture into more distant semantic regions, whereas exploitation requires sustained but linearly scaling cognitive control.

However, contrary to our predictions, CD did not significantly predict inter-response semantic distance in any condition. CD thus appears to track the temporal effort invested in search rather than the semantic character of retrieved ideas, potentially indexing moments of heightened engagement or release from fixation (Critchley et al., 2001). Therefore, CD may track ideation difficulty and level of task engagement (Critchley & Harrison, 2013; Jennings & van der Molen, 2005) rather than the magnitude of conceptual jumps achieved. In optimal foraging terms, CD indexes the energetic investment in search rather than the spatial displacement obtained (Charnov, 1976; Pyke et al., 1977). The nonlinear amplification during exploration aligns with foraging models predicting escalating costs as animals deplete local resource patches and must travel increasing distances to find new ones (Hills et al., 2012). Moreover, each additional millisecond of heart rate slowing during exploratory search produced disproportionately longer thinking times, consistent with the computational demands of retrieving remote associations from sparsely connected semantic network regions (Kenett et al., 2014; Benedek et al., 2017).

An alternative interpretation is that CD may partially reflect motor preparation preceding verbal output (Obrist, 1968); however, supplementary analyses that modelled button-press events separately revealed dissociable patterns: motor-related activity was characterized by beta-band modulation and cardiac acceleration consistent with parasympathetic withdrawal during action execution (Skora et al., 2022; Hashemi et al., 2019), whereas ideation-related activity showed alpha desynchronization coupled with cardiac deceleration. The distinct frequency profiles and opposite cardiac directionalities indicate that the observed CD-alpha coupling reflects ideation-related processes rather than motor confounds.

Although CD did not predict the inter-response semantic distance of generated ideas, it was associated with neural signatures of exploratory processing. Greater cardiac slowing during spontaneous exploration co-occurred with alpha desynchronization over posterior-occipital regions, consistent with angular gyrus engagement during creative semantic restructuring (Fink & Benedek, 2014; Thakral et al., 2020). The angular gyrus, a posterior DMN hub activated during divergent thinking (Beaty et al., 2014, 2016), coordinates retrieval from distributed semantic networks and undergoes functional reorganization during creative ideation (Ding et al., 2025). Converging causal evidence further highlights this role, where disrupting angular gyrus activity via repetitive transcranial magnetic stimulation leads to paradoxical increases in alpha power alongside impaired performance on a semantic decision task (Capotoso et al., 2017). More broadly, posterior alpha desynchronization has been linked to reduced sensory gating and enhanced semantic processing (Klimesch et al., 1998; Jensen & Mazaheri, 2010). Together, the coupling of cardiac deceleration and posterior alpha desynchronization suggests that visceral signals may align with neural states that facilitate exploratory semantic search. Rather than directly shaping the outcome of idea generation, cardiac deceleration may reflect a process-oriented mechanism.

Convolution modelling provides strong support that these effects reflect ideation-related rather than motor-related processes. This analytic approach was advantageous for disambiguating cognitive from motor and physiological contributions to observed neural activity, due to its ability to separate temporally overlapping responses and estimate temporo-spectral impulse response functions for continuous stimuli (Litvak et al., 2013). Traditional event-related designs time-lock analyses to discrete events (e.g., stimulus onset, response), assuming neural activity during these windows reflects only the cognitive process of interest. However, creative ideation involves continuous, self-paced cognitive engagement punctuated by discrete motor responses (button presses to indicate response), creating temporal overlap between ideation-related neural activity, motor preparation, and motor execution (Litvak et al., 2013). Moreover, cardiac activity generates electrical field artifacts in scalp EEG recordings that can spuriously correlate with behavioral measures when heart rate varies systematically with cognitive load (Dirlich et al., 1997; Kwon et al., 2020). Convolution modeling addresses these confounds by simultaneously estimating separate regressors for ideation behavior and button-press events, while accounting for artifacts such as movement or cardiac signals, thereby partitioning variance attributable to each process (Auksztulewicz et al., 2023; Hein et al., 2023; Lai et al., 2024).

Together, these findings support a mechanistic account in which parasympathetically mediated cardiac deceleration modulates the temporal dynamics of ideational engagement during creative cognition. They reinforce emerging views that cognition relies on embodied, process-sensitive mechanisms that extend beyond cortical computations alone (Allen et al., 2019; Babo-Rebelo et al., 2016; Park & Tallon-Baudry, 2014). These findings suggest that cardiac deceleration is as a process-sensitive interoceptive marker that tracks the temporal effort of semantic search during creative ideation, functionally dissociable from both the content of ideas generated and the motor acts through which ideas are expressed.

### Cardiac phase modulates ideational strategies, not contents

We initially hypothesized that creative exploration, both spontaneous and directed, would preferentially occur during the systolic phase of the cardiac cycle, given evidence that systolic baroreceptor firing transiently attenuates exteroceptive processing and may facilitate internally oriented cognition (Edwards et al., 2007; Garfinkel et al., 2014; Park & Blanke, 2019). However, the present findings revealed the opposite pattern. Individuals exhibiting greater diastolic response bias demonstrated greater spontaneous inter-response semantic distances and higher accuracy during directed-switching, whereas individuals exhibiting greater systolic response bias demonstrated lower inter-response semantic distances and directed-persistence failures, even after controlling for baseline heart rate differences. These results suggest a more nuanced interpretation of cardiac afferent signaling.

During systole, baroreceptor discharge projects to nucleus tractus solitarius and downstream cortical targets including insular and anterior cingulate regions (Allen et al., 2019; Critchley & Harrison, 2013; Park & Tallon-Baudry, 2014). Although such signaling may reduce exteroceptive interference, it may simultaneously reduce cortical excitability and response flexibility (Garfinkel et al., 2014; Azzalini et al., 2019). By contrast, diastole represents a relative quiescent afferent window in which baroreceptor activity is minimal. Reduced baroreceptor input during diastole may permit broader associative activation within semantic networks, potentially facilitating the flexible recombination processes required during idea generation (Benedek et al., 2016; Fink & Benedek, 2014). In this framework, diastolic timing appears to be biased towards exploratory cognitive processing, whereas systolic timing biases cognition toward exploitative processing.

Importantly, phase effects were robust at the level of individual differences and condition-averaged tendencies rather than as causal trial-by-trial differences. Rayleigh tests showed no consistent group-level cardiac phase clustering, and mixed-effects analyses revealed limited and context-specific trial-level associations. This pattern is consistent with models in which interoceptive signals bias cognitive operations probabilistically rather than deterministically triggering discrete cognitive transitions (Allen et al., 2019; Seth & Friston, 2016). The D-AUT condition further clarifies the interaction between physiological timing and task constraints. When exploration was externally directed, successful switching was associated with reduced systolic dominance relative to switching failures. Conversely, directed exploitation showed weaker and less consistent phase effects. Such context sensitivity suggests that cardiac phase does not exert uniform directional influence but may interact with control demands and goal states – an interpretation supported by emerging evidence that cardiac timing modulates perceptual and cognitive processes in task-dependent ways (Garfinkel et al., 2014; Motyka et al., 2019).

Together, these findings potentially position cardiac phase as a probabilistic modulator of creative behavior. Rather than exerting momentary control over idea generation, interoceptive timing appears to bias semantic search behaviors, extending embodied accounts of cognition into the domain of creative ideation (Park & Tallon-Baudry, 2014; Allen et al., 2019; Babo-Rebelo et al., 2016).

### EEG time-frequency mechanisms of cardiac-ideational coupling

Our initial hypothesis was that cardiac phase effects would manifest through EEG time-frequency signatures associated with internally directed attention and cognitive control. Specifically, we predicted that systolic timing would align with neural markers of exploration, given prior evidence that systole transiently attenuates exteroceptive processing and modulates cortical excitability (Azzalini et al., 2019; Garfinkel et al., 2014; Park et al., 2014). The present findings refined this prediction by demonstrating a more nuanced and condition-specific pattern of cardiac–brain alignment.

The alpha band showed distinct cardiac couplings depending on cortical region: posteriorly, cardiac deceleration predicted alpha desynchronization during semantic exploration; frontally, diastolic response tendency predicted frontal alpha synchronization. The posterior pattern reflects enhanced semantic processing and reduced sensory gating (Klimesch et al., 1998; Jensen & Mazaheri, 2010), consistent with the angular gyrus engagement already described. The frontal pattern is consistent with alpha’s role in gating exteroceptive interference to allow internally directed associative processing during creative ideation (Agnoli et al., 2020; Benedek et al., 2016; Luft et al., 2018). This regional dissociation – posterior alpha desynchronization coupled with frontal alpha synchronization – suggests distinct functional roles: posterior regions reflect semantic retrieval demands while frontal regions reflect internal attentional orientation. Exploratory analysis revealed similar patterns for originality-related alpha desynchronization in posterior regions.

Theta band activity revealed a systematic relationship between systolic cardiac bias and cognitive control mechanisms. Systolic response bias predicted elevated frontal theta amplitudes during spontaneous ideation related to both inter-response semantic distance and originality. Individuals who preferentially responded during systole, and who exhibited a preference for spontaneous semantic exploitation, showed stronger frontal theta, consistent with enhanced mental set maintenance and reduced cognitive flexibility (Cavanagh & Frank, 2014). Frontal midline theta has been attributed to generators in the anterior cingulate cortex, prefrontal cortex, and hippocampus (Cohen, 2011; Fujisawa & Buzsáki, 2011), and increases during tasks requiring sustained attention, working memory maintenance, and top-down control (Cavanagh et al., 2012; Enriquez-Geppert et al., 2014; Scheeringa et al., 2009). The theta-systole coupling suggests that individuals prone to responding during heightened baroreceptor signaling recruit greater cognitive control mechanisms, potentially compensating for physiological constraints on exploratory switching or reflecting trait-level tendencies toward controlled, exploitative processing. This interpretation aligns with evidence that the cardiac cycle modulates frontal theta in task-dependent and response-congruent manners (Adelhöfer et al., 2020).

Finally, beta band activity showed associations with both systolic cardiac bias and task-specific control demands. During spontaneous ideation, systolic response tendency predicted beta synchronization in anterio-frontal and parieto-occipital regions, further supporting the link between systolic timing and sustained attentional control. Beta oscillations are implicated in maintaining the current cognitive or motor status quo (Engel & Fries, 2010; Jenkinson & Brown, 2011), with synchronization reflecting the stabilization of task-relevant representational states. During directed-persisting, exploratory analyses suggested trends for beta synchronization in parieto-temporal and parieto-occipital regions related to accurate categorical persistence, though these effects were weak. The convergence of beta effects with systolic timing and exploitative strategies is consistent with beta synchronization indexing the neural stabilization required for semantic persistence, though given the exploratory nature of these findings, this interpretation requires caution.

These findings partially support our EEG time-frequency hypotheses while revealing important constraints. We predicted alpha synchronization would be most notable in right temporal regions reflecting interoceptive processes (Ceunen et al., 2016; Chong et al., 2017) yet observed that the strongest cardiac-alpha coupling appeared in frontal regions, with posterior effects specific to cardiac deceleration rather than phase. This regional specificity may reflect thalamic relay of nucleus tractus solitarius projections to frontal cortical regions (Critchley & Harrison, 2013), though the specific anatomical pathway remains to be characterized. Additionally, it is possible that cardiac signals may modulate alpha power through heartbeat-evoked potentials (Villena-González et al., 2017) and heart rate variability correlations (Luft & Bhattacharya, 2015; Magosso et al., 2019; Zaccaro et al., 2018), with these effects potentially exhibiting regional variability that warrants systematic investigation in future work.

These time-frequency findings extend previous findings of cardiac-phase modulation in perception and decision-making (Garfinkel et al., 2014; Park et al., 2014) to the domain of semantic search, situating creative ideation within a broader framework of cardiac–cortical coordination. Crucially, this coordination is not uniform but specifically differentiated: alpha couples selectively with cardiac variables to support semantic retrieval and internal attentional orientation, theta links systolic dominance to cognitive control and categorical persistence, and beta synchronization aligns with exploitative set-maintenance. Together, these findings demonstrate that the body’s rhythmic influence on creative cognition operates through frequency-specific oscillatory channels, each serving a distinct function in the temporal organization of ideational search.

### Interoceptive rhythms as embodied constraints on semantic foraging

These findings converge with broader theoretical frameworks positioning creative cognition as embodied semantic foraging emerging from multi-scale interactions between physiological, neural, and cognitive processes. Optimal foraging theory presents creative ideation as analogous to animals searching for resources in physical space (Hills et al., 2012, 2015; Todd & Hills, 2020): ideation alternates between local exploitation (area-restricted search within semantic categories) and global exploration (long-distance jumps to distant regions). In this model, transitions are governed by diminishing returns within depleted patches (Charnov, 1976): the marginal value theorem predicts patch-leaving when local rates fall below global averages (Stephens & Krebs, 1986). Computational models of semantic search confirm these foraging dynamics, with individuals navigating semantic networks through similarity-driven local search and frequency-modulated global jumps (Hills et al., 2012; Abbott et al., 2015; Kumar et al., 2024).

The present results demonstrate that semantic foraging operates under embodied physiological constraints. Cardiac deceleration tracks the escalating effort of exploratory search, analogous to energetic costs of long-distance spatial movements in animal foraging (Pyke et al., 1977). Cardiac phases bias when transitions occur, providing rhythmic temporal windows favoring exploration (diastole) or persistence (systole), akin to the way circadian or ultradian rhythms modulate foraging patterns in animals (Daan & Aschoff, 1982). Individual differences in cardiac-cortical coupling predict inter-subject variation in explore-exploit tendencies, paralleling individual differences in animal foraging strategies shaped by metabolic state and environmental history (Stamps, 2016). This positions creative ideation as fundamentally embodied: visceral signals continuously bias the cognitive operations underlying semantic search as functional constraints shaping when and how ideas emerge.

The salience network provides a plausible integrative substrate for these interactions. The anterior cingulate and insula, the core hubs of the salience network, activate during creative tasks (Beaty et al., 2015; Goulden et al., 2014) and orchestrate dynamic shifts between DMN-driven associative exploration and ECN-driven controlled exploitation (Menon & Uddin, 2010; Sridharan et al., 2008). These same regions receive primary interoceptive inputs (Craig, 2009), generate bodily prediction errors (Seth & Friston, 2016), and modulate autonomic output (Critchley & Harrison, 2013). Within this architecture, cardiac rhythms may contribute temporal structure that biases how and when cortical networks engage in exploration versus exploitation, without affecting the creative quality of generated ideas.

Together, these findings suggest that creativity emerges from dynamic brain–body coordination rather than from cortical processes in isolation. Across all three levels of analysis – cardiac deceleration, cardiac phase and cardiac-brain coupling – the data converge on a single principle: interoceptive rhythms probabilistically shape the temporal and strategic organization of ideational search without determining its semantic content. Semantic foraging is thus embedded within rhythmic interoceptive cycles that constrain the when and how of idea generation, aligning creative cognition with broader neurovisceral models of embodied decision-making and adaptive behavior (Thayer & Lane, 2009; Park & Tallon-Baudry, 2014). Rather than reducing creativity to cardiac dynamics, this framework situates creative ideation within a hierarchically organized system in which physiological rhythms and cortical activity jointly constrain semantic search.

### Limitations and future directions

Several limitations warrant consideration. First, we did not assess potential confounds including exercise habits, body mass index, or respiratory activity, which may influence cardiac dynamics and modulate cardiac-creativity relationships (Grossman & Taylor, 2007; Rominger et al., 2022). Physical fitness enhances heart rate variability and may strengthen cardiac-cortical coupling (Wallman-Jones et al., 2021), potentially contributing to the individual differences we observed. Second, the experimental design randomized directed exploration and directed exploitation trials rather than presenting them in separate task blocks. This approach was chosen to minimize anticipatory responses, but it may have attenuated condition-specific effects: repetitive task switching is known to impact creative performance (Lu et al., 2017). We addressed this statistically by modelling trial number and intra-individual variability using mixed-effects approaches, but future work using blocked designs could establish whether stronger cardiac-ideational coupling emerges with sustained task engagement in a single strategy mode. Third, cardiac phase interpretations require caution given natural probability asymmetries: diastole lasts longer than systole, with this ratio varying as a function of heart rate (Fridericia, 2003). We controlled for inter-beat interval length and tested the interaction between cardiac phase and heart rate, but our results showed no significant interactions, indicating that cardiac phase was a robust predictor of ideation behavior. We also used circular statistics to account for individual variability in cardiac phase probabilities. However, the relatively short R-R intervals (∼800 ms) may introduce temporal uncertainty when aligning self-paced responses to cardiac phases. Traditional cardiac phase paradigms time-lock external stimuli to specific phases, providing precise phase control; our self-paced design sacrifices this precision for ecological validity. Future work incorporating externally timed prompts could establish stronger trial-level phase effects at the cost of limiting the ecological validity of the ideal paradigm. Fourth, our participants were not instructed to use a specific response hand, limiting hemispheric dissociation of motor-related spectral activity from ideation-related processes. While convolution modelling isolated button-press signals, hand-specific analysis could better establish lateralized effects. Fifth, we did not focus our analyses on upper and lower alpha bands as in Mastria et al. (2021); instead, we focused on broader alpha, beta, and theta bands, based on prior evidence that visceral signals preferentially modulate these frequency ranges (Kern et al., 2013; Luft & Bhattacharya, 2015). Sixth, spontaneous ideation effects emerged primarily at the participant level while directed condition effects appeared at the trial level, potentially a reflection of our study design, in which the spontaneous trials were assessed as a whole, while the directed trials were separated into directed-switch and directed-persist. Additionally, it may reflect greater inter-individual variability in spontaneous strategy selection or reduced sensitivity given high accuracy in directed condition (directed-switch: 85%, directed-persist: 68%). We addressed this in part by including inter-response semantic distance as a continuous measure of ideational breadth and by applying trial-level mixed-effect models. Finally, our correlational design precludes causal claims regarding directional cardiac-ideational relationships. While we controlled for motor confounds and employed temporal precedence where possible (cardiac deceleration preceding responses), true causal inference requires experimental manipulation or time-lagged analyses. Our convolution modelling averages within conditions rather than preserving single-trial temporal dynamics, preventing trial-by-trial lagged regression for EEG activity (Litvak et al., 2013). Future work combining high-temporal-resolution cardiac metrics with single-trial oscillatory analyses could establish whether cardiac changes predict subsequent ideational shifts or vice versa, potentially revealing bidirectional feedback wherein ideational strategy selection modulates cardiac state which in turn influences subsequent ideation.

### Conclusions

This study suggests that creative ideation unfolds within temporally structured, embodied interactions between cardiac signals and cortical activity. Cardiac deceleration predicted response time in a condition-specific manner, scaling nonlinearly during exploratory search and linearly during exploitative search; cardiac phase biased ideation strategies, with diastolic timing linked to exploration and systolic timing linked to exploitation; and individual differences in cardiac-cortical coupling predicted variation in strategy selection. These findings extend prevailing cortico-centric accounts of creativity by suggesting that semantic search is embedded within interoceptive rhythms. Rather than operating independently of bodily state, idea generation appears coordinated within ongoing neurovisceral cycles. In this view, creative thinking reflects dynamic brain-body integration, in which interoceptive timing and cortical activity jointly constrain semantic exploration and exploitation. By situating creative cognition within a multi-scale framework linking physiological rhythms, oscillatory coordination, and semantic search dynamics (D’Angelo & Jirsa, 2022; Kotler et al., 2025; Song et al., 2025), this study identifies cardiac–cortical alignment as a previously underexamined dimension of creative thought. As models of higher cognition increasingly incorporate embodied and network-level perspectives, understanding how visceral rhythms interact with cortical computations may prove essential for explaining individual differences in the balance between exploration and exploitation during human creative ideation.

## Methods

### Participants

Fifty-nine participants (39 female; age range: 19-42 years, M = 23, SD = 4.76) participated in the study. All were right-handed, either native or proficient English speakers and reported no history of neurological or psychiatric disorders. Three participants were excluded due to either failure to understand task instructions or loss of more than 50% of cardiac or neural data. The sample size was informed by previous studies on cardiac cycle effects in cognition (e.g., Garfinkel et al., 2014) and studies on switching and persisting during creativity using EEG (e.g., Mastria et al., 2021). All participants provided signed informed consent before participation. Participants either received course credits or £25 for their participations. The study followed the Declaration of Helsinki and was approved by the Ethics Committee of the Department of Psychology at Goldsmiths University of London.

### Task Design

The study employed a within-subjects design to examine the influence of cardiac and neural dynamics on creative semantic switching. Two versions of the Alternate Uses Task, AUT (Guilford, 1967) were administered: a spontaneous condition, which allowed unrestricted idea generation, and a directed condition, which imposed semantic constraints on semantic switching. This structure enabled dissociation between internally generated (spontaneous) and externally cued (directed) ideational transitions.

In the spontaneous condition, the standard AUT, participants generated alternative uses for everyday objects without restriction on strategy, assessing natural fluctuations between exploration and exploitation. In the directed condition, each trial specified whether participants should remain within the same semantic category (persist) or deliberately switch to a new one (switch) upon generating a new idea. The D-AUT was designed to probe controlled transitions in semantic retrieval and was informed by previous work on semantic search dynamics and executive guidance of creative thought (Wu & Koutstaal, 2022).

The two task conditions were matched in duration and response structure, enabling comparison of ideational dynamics under free and constrained conditions. Both behavioral responses (button press, ideation output) and physiological signals (EEG, ECG) were time-locked for trial-level convolution analysis.

### Procedure

All participants were tested individually in a sound-attenuated, dimly lit room. Upon arrival, they were briefed on the general aims of the study and fitted with a 64-channel EEG cap and ECG electrodes. Following impedance checks and signal verification, participants completed a short practice session to familiarize themselves with the task format and response protocol.

The experiment comprised two conditions of the Alternate Uses Task (AUT): spontaneous and directed. In the spontaneous condition, participants were presented with a common object (e.g., brick, shoe) and instructed to generate as many alternative, uncommon uses as possible within 2 minutes, without specific constraints. Following a break, participants were instructed to complete the D-AUT. In the directed condition, participants were presented with a cue word and instructed either to persist within the current category or to switch to a new semantic category based upon their preceding response. Instructions were clearly displayed on screen at the start of each trial (see *Supplementary Task Stimuli* for specific instructions and cue words).

Participants indicated the onset of each idea by pressing a response button with their dominant hand, immediately followed by a verbal report of the idea. Verbal responses were recorded using a table-mounted microphone and transcribed for later semantic analysis. Participants began the experiment completing five practice trials of the standard AUT, with each trial lasting 30 seconds. Then, participants completed 10 spontaneous (AUT) trials, with each trial lasting 2 minutes. For each object, they generated as many creative uses as they could. After a short break, participants then completed 10 directed trials. For the D-AUT, within each trial (i.e., across responses for each object) response directions varied randomly between switch and persist according to a pre-defined order that did not vary across participants. Optional breaks were provided between trials to minimize fatigue.

Throughout both tasks, EEG and ECG signals were continuously recorded. The timing of button presses, and verbal responses were logged to enable alignment with physiological data. Participants completed the AUT and the D-AUT in addition to other tasks not discussed in this paper. In total, sessions lasted around two-and-a-half hours per participant.

### EEG and ECG Acquisition

Continuous EEG and ECG data were recorded simultaneously using a BioSemi® ActiveTwo systems with 64 Ag/AgCl scalp electrodes placed according to the international 10-20 system. The vertical and horizontal eye-movements (EOG) were monitored by electrodes above and below the right eye and from the outer canthi of both eyes, respectively. Additional external electrodes were placed on both left and right ear lobes as reference. The ECG was recorded via two flat electrodes placed two inches below the right clavicle and two inches above and three across (medially) the left hipbone using the two remaining external channels with a bipolar ECG lead II configuration. The sampling frequency was 512 Hz.

#### EEG Pre-processing

EEG data were processed using the EEGLAB Toolbox (Delorme & Makeig, 2004). Upon import, electrode positions were aligned using the ‘standard_1005.elc’ (MNI configuration). The EEG data were re-referenced to the average of the two earlobes electrodes and then high-pass filtered at 0.5 Hz, low-pass filtered at 100 Hz, and notch filtered at 50 Hz (48-52 Hz). Behavioral event markers (button presses for “speak start” and “speak end”) from PsychoPy were used to time-lock EEG data. Epochs were extracted from - 3 s to +2 s around the “speak start” marker. The data were inspected visually for artefacts, and noisy channels or segments were removed and interpolated (3 or fewer channels removed). Epochs with response times below 3 s were excluded from further analysis. Independent components analysis (ICA, using the Infomax algorithm) was run to remove stereotyped artefacts, including blinks and cardiac field artifacts. These components were identified using visual inspection of time series, topographies and spectral characteristics and removed prior to reconstruction of the cleaned EEG signal.

#### ECG Pre-processing

The ECGdeli toolbox (Pilia et al., 2021) was used to process ECG signals. Continuous ECG data were first baseline-corrected and then filtered with a 0.3 Hz high-pass and a 120 Hz low-pass filter (based on Pilia et al., 2021). Line noise was further removed using an isoline correction procedure. Waveform delineation was automatically performed to extract timestamps for P-wave, QRS complex, and T-wave, followed by visual inspection to verify accuracy.

We opted for an individualized approach which considered between and within-subjects variations. Specifically, we defined systole as the interval from the R-peak to the T-wave offset, and diastole as the interval from the T-wave offset to the next R-peak (Bury et al., 2019; Garfinkel & Critchley, 2016; Kunzendorf et al., 2019). A binary cardiac afferent variable was created for each behavioral event, indicating whether it occurred during systole (1) or diastole (0).

Then, we computed the cardiac phase angle at each response onset by locating its relative position between the preceding and following R-peaks and mapping this interval to a 0-2π radian scale (Kunzendorf et al., 2019). These values were used for circular statistical analysis of cardiac phase bias. Cardiac activity was quantified by computing inter-beat intervals (IBIs) across trials. Mean IBIs were derived from three consecutive intervals as follows: 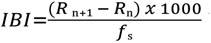 where *R*ₙ and *R*_n₊₁_ are successive R-peaks, *fₛ* is the sampling rate in Hz. Cardiac deceleration (CD) was calculated using a moving average of six consecutive IBIs: two preceding, one concurrent with, and three following each behavioral response (Motyka et al., 2019), providing an index of vagal engagement linked to ideation timing.

#### Time-Frequency Decomposition and Convolution Modelling

To assess how ideation behaviors modulated time-frequency (TF) EEG activity, we used convolution modelling of time-frequency (TF) data (Litvak et al. 2013). This method provided estimates of TF amplitude changes related to a series of discrete and parametric event-related regressors. Our convolution GLM analyses were conducted using SPM12 and FieldTrip, and followed the protocol established in Hein and Herrojo Ruiz (2022)

Preprocessed EEG data were first converted to the SPM format, followed by TF decomposition using a Morlet wavelet transform (5 cycles) from 4 to 30 Hz. For each subject, we constructed design matrices containing event-locked regressors. These event-related design matrices were constructed using task-specific regressors. For the AUT, two types of regressors were modelled: (i) button presses (discrete), and (ii) semantic distance (parametric continuous regressors). For the D-AUT, five discrete regressors were defined: (i) switch_hit: participants were cued to switch and did switch appropriately, (ii) switch_miss: participants were cued to switch but persisted instead, (iii) persist_hit: participants were cued to persist and did persist appropriately, (iv) persist_miss: participants were cued to persist but switched instead, and (v) button-presses (to capture motor-related activity). A third convolution GLM was conducted for exploratory testing on originality ratings. This was done using AUT (i) button presses (discrete), and (ii) originality (continuous) as regressors. Originality was operationalized using semantic distance between each response and the cue word, while switching scores were calculated based on the semantic distance from the immediately prior response. Stimulus presentation was not explicitly modelled due to the structure of response sets and the exclusion of first responses from analysis (where there is no “switching/persisting” from cue to response).

Each regressor was convolved with a 20th-order Fourier basis set (40 basis functions: 20 sines and 20 cosines), allowing the modelling of spectral modulations up to ∼9 Hz across an 8 s window (-5 to + 3 s around event onset). This configuration ensured sensitivity to both transient and sustained oscillatory effects linked to ideation, semantic switching, and motor responses. Models were estimated separately for each condition, using SPM12 and custom MATLAB scripts. Resultant time-frequency images (in arbitrary units) were stored in 3-D image format (electrode x time x frequency), preserving the spectro-temporal profiles of task-related dynamics. For group-level analysis, mean time-frequency amplitudes were extracted from three frequency bands: theta (4 – 6 Hz), alpha (8 – 12 Hz), and beta (13 – 30 Hz); averaging was performed over a pre-event window of interest (-2 to 0 s) for each regressor at each electrode. Sensor-level data were grouped into nine bilateral regions of interest (ROIs): anterior frontal (Fp1, AF3, AF7), frontal (F3, F7), fronto-central (FC1, FC3, FC5, FT7, FT9), centro-temporal (T7, C5, C3, C1), centro-parietal (CP1, CP3, CP5, TP7, TP9), parieto-temporal (P1, P3, P5, P7, P9), parieto-occipital (P03, P07, P09), occipital (O1, O2), and the right-hemisphere homologues. Midline electrodes (Fz, FCz, Cz, CPz, Pz) were included in the analyses as a separate set due to their important in ideation-related processes and to allow testing for hemispheric differences (Mastria et al., 2021).

The resulting time-frequency images were converted to FieldTrip-compatible structures (Oostenveld et al., 2011) for statistical analyses. Importantly, unlike traditional time-frequency approaches, no baseline correction was applied, as GLM-based modelling incorporates baseline dynamics by accounting for event latency variability and overlapping temporal responses (Litvak et al., 2013). Motor-related beta rebound was reliably captured by discrete button-press regressors, serving as a control for action-related oscillatory dynamics (see Supplementary Figure 2). This approach allowed us to isolate ideation- and originality-related modulations beyond sensorimotor confounds.

#### Behavioral Scoring

To quantify ideational performance across conditions, we employed multiple scoring approaches for responses generated in the AUT and the D-AUT.

#### Behavioral Ratings

Semantic distance (SemDis: Beaty & Johnson, 2021) was used as a continuous, trial-level proxy for exploration and exploitation in idea generation. SemDis quantifies the semantic distance between a given response and its immediate predecessor, operationalizing exploration as higher inter-response semantic distance. Only in the case of the D-AUT, SemDis scores were z-scored across participants to enable standardized classification. Responses with z-scored SemDis values above the individual mean were designated as “switches” (exploratory), whereas those below were designated as “persist” (exploitative). These were then used to assess if the directed trial was a “hit” or a “miss”.

To independently validate this computational classification, two external raters (blind to the study’s aims) were recruited and trained to classify responses manually as a switch or persist from the response prior, creating a binary marker for ideation behaviour. Inter-rater agreement was moderate (Cohen’s *κ* = .64, *p* < .001). Agreement between automated SemDis-based classifications and human ratings was quantified using binary logistic regression. For the first rater, the model was statistically significant, *χ²* (1) = 665, *p* < .001, explaining 15% of the variance (Nagelkerke *R*²); for the second rater, *χ*² (1) = 465, *p* < .001, explaining 13% of the variance. These results provided internal validation of the semantic classification procedure used to derive ideation modes across the experiment.

### Statistical Analyses

All statistical analyses were performed using R (versions 2024 and 2025), incorporating packages *lme4, lmerTest, afex,* and *circular.* Analyses were structured to evaluate how cardiac indices – inter-beat interval (IBI), cardiac deceleration (CD), and cardiac phase – modulated three key behavioral outcomes: (i) response time, (ii) semantic distance, and (iii) accuracy under directed-switch or directed-persist constraints. Four complementary levels of analysis were employed: (i) Individual differences to identify stable cardiac influences on ideation style, (ii) Trial-level mixed-effects models to capture moment-to-moment cardiac-behaviour coupling, (iii) Simple linear regressions linking cardiac measures with task-evoked EEG amplitude changes across canonical frequency bands, and (iv) Circular statistics to model the cyclical nature of cardiac phases. Finally, we conducted exploratory analyses related to possible motor confounds in the beta frequency band as well as exploratory analyses related to the originality of responses (semantic distance from the cue word to the response).

#### Individual Differences Analysis

To examine between-participant variation in the relationship between cardiac phase and ideation behavior, we conducted linear regressions using participant-averaged data. Binary trait-level variables were transformed into proportions (i.e., proportion of ideations during systole vs. diastole) to permit continuous modelling. To account for potential confounds, follow-up multiple linear regression models included average heart rate and response time as covariates. Model assumptions (normality, homoscedasticity, and multicollinearity) were verified, and variance inflation scores indicated no multicollinearity.

A two-way ANOVA was conducted to assess whether cardiac phase (indexed as the proportion of systolic responses) varied by condition (directed-switch vs. directed-persist) and ideation behaviour (switch vs. persist). Post-hoc contrasts were performed using estimated marginal means with Holm correction for multiple comparisons.

For EEG data, we focused on within-condition relationships (i.e., within AUT and within D-AUT), rather than direct comparisons between spontaneous and directed tasks. Pre-processed EEG band power (theta, alpha, beta) and cardiac indices (cardiac deceleration and cardiac phase) were merged at the participant level for those with valid data across both modalities. EEG power values were first structured in long format, including factors for frequency band, region of interest (ROI), and hemisphere. Repeated-measures ANOVA with with Greenhouse-Geisser correction revealed no significant hemispheric differences for either condition, so data were collapsed across hemispheres.

Descriptive statistics (participant count, trial number, mean power, and mean cardiac metrics) were computed for each band x ROI combination. EEG power was regressed onto cardiac predictors: mean cardiac deceleration at response and mean systolic/diastolic phase at response, separately for each frequency band and ROI. Dependent variables included semantic distance (sem_dis), and originality scores (sem_orig) in the spontaneous condition, and response accuracy (switch_hit, switch_miss, persist_hit, persist_miss) in the directed condition. Predictors were tested independently, and *p*-values were adjusted for multiple comparisons using Benjamini-Hochberg false-discovery rate (FDR).

To explore the general inhibition hypothesis posed by Obrist (1968) which proposed that cardiac deceleration reflects reduced peripheral muscular activity during action execution rather than attentional modulation as argued by Lacey and Lacey (1970), we also included frequency amplitudes time-locked to button press as a supplementary analysis. This allowed us to test whether cardiac-brain coupling reflected motor output suppression versus cognitive reallocation.

#### Circular Analysis

Given the inherently cyclical nature of cardiac activity, we employed circular statistics to assess the timing of responses across the cardiac cycle. Response timing was converted into radians (0-2π), and Rayleigh’s test was used to assess non-uniformity of response distributions within each condition (spontaneous, directed-switch/persist). For each participant, the mean cardiac phase (*μ*) and resultant vector length (*r*) were calculated to index preferred response phase and concentration, respectively.

To examine whether participants’ mean response phase were associated with their mean SemDis scores, we applied a vector-based method, of circular-linear correlations (Kunzendorf et al., 2019), with significance evaluated via *χ*² approximation and 10,000 random permutations (r₍cl₎ = √[cor(SemDis, sinμ)² + cor(SemDis, cosμ)²]). Correlations between *r* and SemDis were also computed to assess whether greater phase-locking was associated with increased semantic distance. These analyses were repeated for EEG-cardiac phase coupling (*μ* with band power; *r* with band power), within both spontaneous and directed conditions; FDR correction was applied within each task and regressor family to control for multiple comparisons.

#### Mixed-Effects Modelling

Trial-level linear and logistic mixed-effects models were fitted using the lmer() and glmer() functions in R, incorporating random intercepts for participant and trial to account for nested dependencies. Models were constructed separately for each behavioral outcome: (i) spontaneous ideation (semantic distance) - predicted by trial-level CD, IBI, and cardiac phase, (ii) directed-switch accuracy– modelled similarly, and with binary logistic regressions, (iii) directed-persist accuracy – modelled similarly, and with binary logistic regressions, and (iv) response times for each condition – included as predictors and outcomes to test for bidirectional cardiac-behavioral relationships. Cardiac predictors were entered hierarchically, in ascending order of their zero-order correlation coefficients with the dependent variable. Statistical significance was evaluated using Satterthwaite’s approximations for degrees of freedom, and model fits were assessed using conditional and marginal R² values. Logistic regression models for binary outcomes were estimated using the logit link function. For all models that tested cardiac phases as fixed factors, we tested IBI as a covariate because the probability of being in systole or diastole changes as a function of heart rate. As heart rate increases, the absolute duration of systole decreases; however, the proportion of the cardiac cycle occupied by systole (relative to diastole) increases (Fridericia, 2003).

