## Supplementary for "Cardiac Timing Biases Creative Exploration and Exploitation"

**Supplementary Materials**

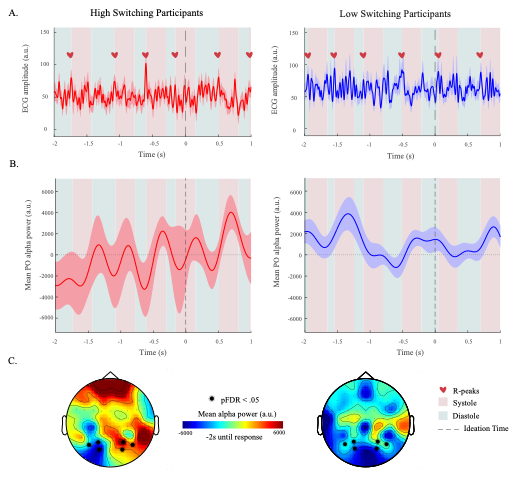

**Supplementary Figure 1. High and low switching during spontaneous ideation.** **A)** Event-locked cardiac time series across the spontaneous (AUT) condition for visualization. Displayed are grand-average waveforms time-locked to response onset (0 s – dotted line), plotted separately for high-switching (blue/right) and low-switching (red/left) participants during the spontaneous (AUT) condition, defined as the top 20 highest and lowest switchers. Shaded error bands denote the standard error of the mean. The heart icons denote approximate R-peak locations identified from local maxima of the averaged trace. Background shading indicates canonical cardiac phases derived from approximate R-peak locations (denoted by a heart icon) in the averaged waveform, with systole defined as the 300 ms interval following each R-peak and diastole defined as the interval from the end of systole to the subsequent R-peak. ECG power is shown in arbitrary units (a.u.), and the time axis spans from 2 s before to 1 s after the response. **B)** Averaged timeseries of parieto-occipital alpha power for high-switching (red/left) and low-switching (blue/right) participants during spontaneous (AUT) condition. The shaded regions represent +/- 1 SEM. An independent-samples t-test showed no significant difference in parieto-occipital (PO) alpha power between high-switching and low-switching participants in the spontaneous condition (sem_prev), *t*(38) = −1.15, *p* = .256, Cohen’s d = −0.36. Alpha power was slightly lower in the high-switching group (*M* = −2435.17, *SD* = 10526.02, *n* = 20) compared to the low-switching group (*M* = 405.94, *SD* = 3282.52, *n* = 20), but this difference was not statistically significant. **C)** Averaged alpha power topology for high-switching (left) and low-switching (right) participants during spontaneous (AUT) condition. Highlighted electrodes represent ROIs with a significant (pFDR < .050) relationship between alpha power and cardiac deceleration.

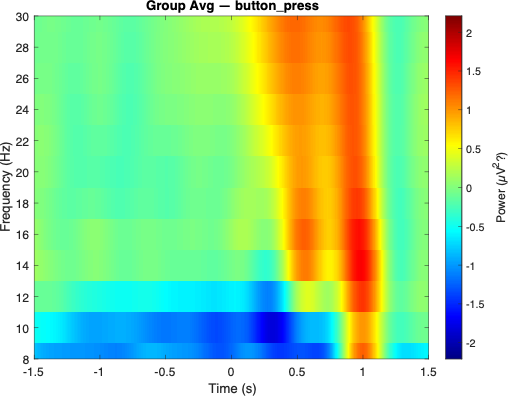

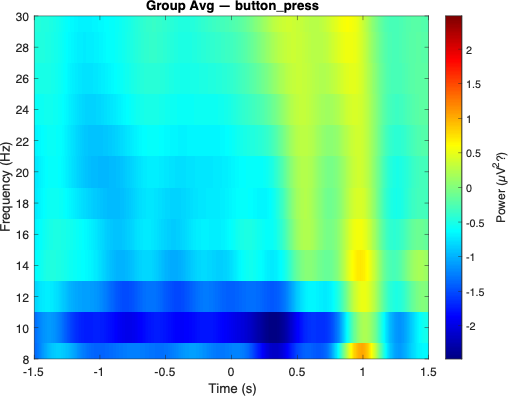

A) Spontaneous Condition

B) Directed Condition

**Supplementary Figure 2. Post-movement beta rebound.** **A)** Time-frequency plots denoting the post-movement beta-rebound in alpha and beta band activity during the: **A)** Spontaneous (AUT) condition timed to the button-press regressor. **B)** Directed (D-AUT) conditions timed to the button-press regressor.

**Supplementary Table 1. Descriptive statistics for Individual Differences**

| Variable | *M* | *SD* |
| --- | --- | --- |
| Average Heart Rate | 76.79 | 8.41 |
| Interoceptive Sensitivity | 82.69 | 14.34 |
| Openness | 28.44 | 2.69 |
| Intellect | 34.51 | 2.87 |
| Creative Achievements | 136.32 | 26.86 |
| Attention Control | 51.75 | 7.77 |

**Supplementary Table 1. Descriptive statistics for individual differences.** This table presents the descriptive statistics for individual differences.

**Supplementary Task Stimuli**

**Alternate uses task cue words**

(1) shoe, (2) fork, (3) belt, (4) newspaper, (5) soap, (6) hat, (7) cardboard, (8) mug, (9) umbrella, (10) balloon, (11) bucket, (12) drinking straw, (13) brick, (14) book, (15) plastic bag, (16) towel, (17) sock, (18) pencil, (19) rope, (20) pillow.

**Alternate uses task instructions**

“For the first task, you will be asked to think of unusual, creative uses for common everyday objects. You will be shown an object and must think of many different creative uses for it. As many different ideas as you can. In the first task there won’t be any additional instructions. The second task is the same, but sometimes you will be asked to think of an idea that is similar to your previous idea, and sometimes you will be asked to think of an idea that is different to your previous idea. You will be shown an object, be tasked to come up with a creative use for it, and then your second idea needs to be either similar or different to your first idea.

Finally, most of these tasks involve thinking of an idea and then saying it out loud. When doing this, make sure to press SPACE when you are ready to say your idea, then say it out loud and clearly, and then press SPACE again. So, in the first task you’ll be shown an object, and you think of an idea for how to use it, you press SPACE, say it out loud, then press SPACE again. Then think of another idea, press space, say it, press space, think of another idea, press space, say it, press space. Make sure you press SPACE and can see the speech bubble before you start speaking.”

**Supplementary Analysis**

**Neural activity during button-press and cardiac deceleration**

As mentioned earlier, we included neural activity from the button-press regressor in our linear testing between amplitudes in the frequency bands and cardiac signals for both the spontaneous condition and the directed conditions. For the spontaneous condition, we found that cardiac deceleration was significantly associated with beta suppression attributed to button-presses in anterio-frontal (β = -19.02, *p* = .010, *pFDR* = .040), centro-parietal (β = -11.66, *p* < .010, *pFDR* = .020), centro-temporal (β = -19.96, *p* < .001, *pFDR* < .001), frontal (β = -24.44, *p* < .010, *pFDR* < .001), fronto-central (β = -24.44, *p* < .001, *pFDR* < .001), and parieto-temporal (β = -9.72, *p* = .010, *pFDR* = .040) regions. There was no relationship between alpha or theta activity during button-press and cardiac deceleration. When analyzing the directed conditions, button-press activity was not significantly associated with cardiac deceleration in any of the regions of interest or frequency band.
